# Promoter activity buffering reduces the fitness cost of misregulation

**DOI:** 10.1101/215186

**Authors:** Miquel Àngel Schikora-Tamarit, Guillem Lopez-Grado i Salinas, Carolina Gonzalez Navasa, Irene Calderon, Xavi Marcos-Fa, Miquel Sas, Lucas Carey

## Abstract

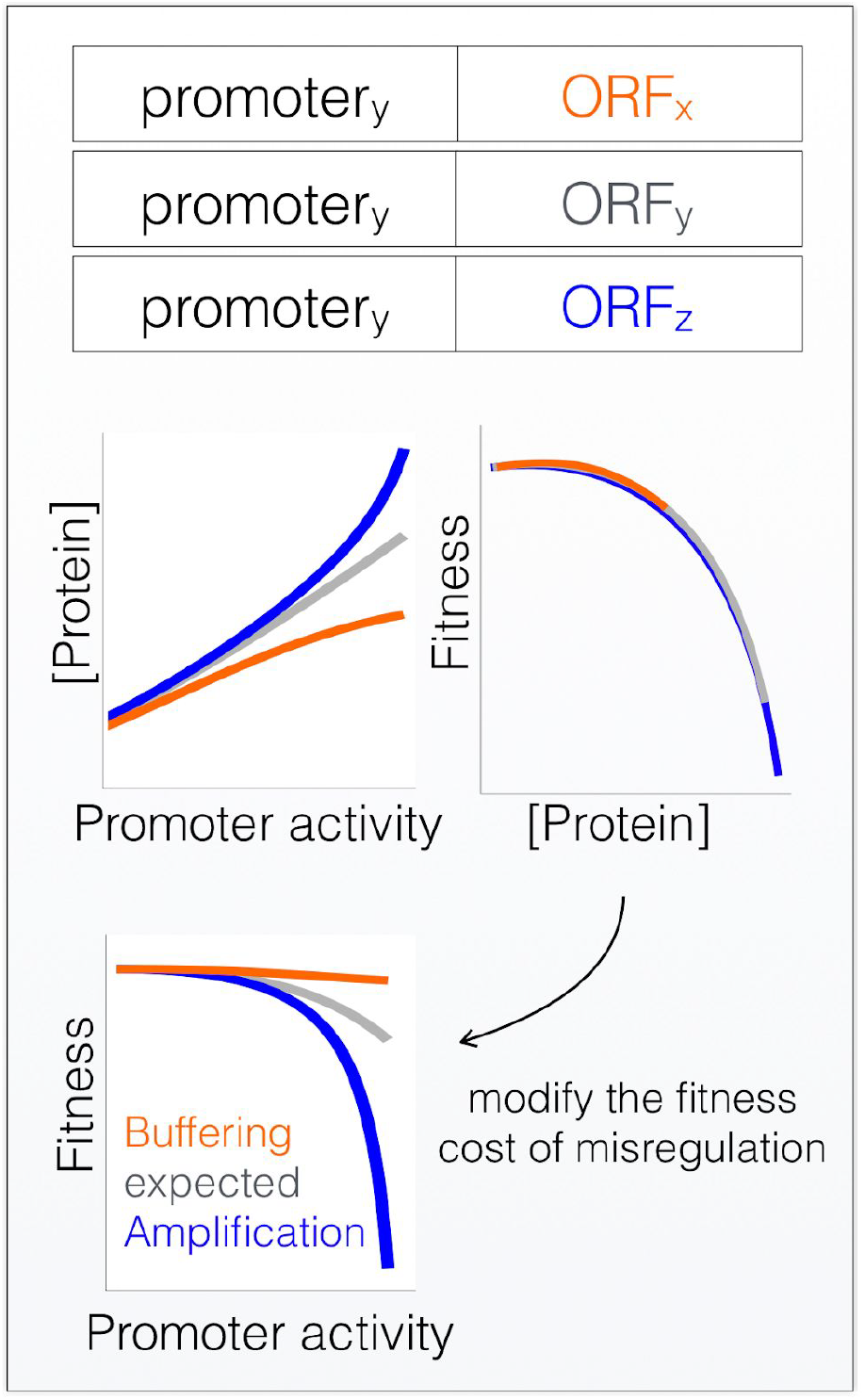

**In Brief:** Systematic promoter replacement reveals that coding sequences and 3’ends of genes play and active role in buffering cells from fitness defects due to misregulation.

**Highlights:**

- The TF dose-response curve -- how gene expression changes as a function of TF concentration -- is encoded throughout the gene, not only in the promoter. Genes with the same promoter have different TF DRCs.
- A coupled experimental system and mathematical model quantifies the intrinsic ability of genes to buffer or amplify promoter activity.
- Promoter activity buffering reduces the effect of misregulation on fitness.

Cells regulate gene expression by changing the concentration and activity of transcription factors (TFs). The response of each gene to changes in TF activity is generally assumed to be encoded in the promoter. Here we show that, even when the promoter itself remains constant, each gene has a unique TF dose response curve. Genes have an intrinsic ability to either buffer or amplify the effects of high promoter activity. We present a coupled mathematical model and experimental system for quantifying this property. Promoter activity buffering can be encoded by sequences in both the open reading frame and 3’end of genes, and can be implemented by both autoregulatory feedback loops and by titration of limiting *trans* regulators. We show experimentally that promoter activity buffering insulates cells from fitness defects due to misexpression. The response of genes to changes in [TF] is encoded by sequences outside of the promoter, and this effect can either insulate or amplify the effects of aneuploidy and misregulation on organismal fitness.

## INTRODUCTION

Understanding how changes in transcription factor activity lead to changes in gene expression is essential for understanding how organisms regulate expression in response to external and internal signals, and for understanding how sequence variation in genomes affects phenotype(Carey et al., 2013; Levo and Segal, 2014; López-Maury et al., 2008; Segal and Widom, 2009). Changes in gene expression are associated with clinically relevant phenotypes such as disease and differential response to drugs, and the current assumption is that much variation in phenotype and expression is due to sequence variation in classical regulatory regions such as enhancers and promoters(Albert and Kruglyak, 2015). However, current mathematical models of gene expression, while good at predicting the effects of small mutations(Levo and Segal, 2014), are less good at the ultimate goal, which is predicting expression from sequence(Karr et al., 2012; Yuan et al., 2007). To quantitatively understand the effects of variation in transcription factor (TF) activity and sequence variation on gene expression, we appear to be missing some key regulatory processes.

Steady state mRNA levels are determined by the balance between synthesis and degradation rates. These rates are unique to each gene, and are determined by the interaction between *cis*-encoded sequence features (e.g., TF binding sites, transcription terminator, translation initiation site context, codon bias, and binding sites for RNA binding proteins (RBPs)) and the concentration and activities of *trans-* factors (e.g., TFs, translation initiation factors, RBPs). Further increasing the complexity of predicting mRNA levels from sequence, translation initiation and elongation both affect mRNA stability(Chen et al., 2017; Dvir et al., 2013; Harigaya and Parker, 2016).

The response of a gene to changes in the concentration and activity of a given *trans* regulator are generally assumed to be encoded in specific regions of a gene. For example, in organisms that lack enhancers, the response of a gene to changes in TF activity are generally assumed to be encoded in the promoter(Ang et al., 2013; Brophy and Voigt, 2014; Carey et al., 2013; Hansen and O’Shea, 2015; Kim and O’Shea, 2008; van Dijk et al., 2015). Individual regulatory elements, such as 5’UTRs, codon usage, and 3’ends (3’UTR + transcription terminator) have been interrogated separately(Chen et al., 2017; Dvir et al., 2013; Radhakrishnan et al., 2016; Shalem et al., 2015, 2013; Yamanishi et al., 2013), with the unstated assumption that there is little or no interaction between distinct regulatory elements. However, recent work suggests the existence of a genetic interaction between promoters and codon usage on steady-state mRNA levels(Espinar et al., 2018), calling into question the idea that these regulatory elements can be fully understood in isolation.

To determine if sequences external to the promoter can influence the response of a gene to changes in [TF] we generated a library of 42 native yeast genes in which each gene is expressed under the control of the same promoter, with the same 5’UTR, and N-terminally tagged with a codon-optimized mCherry. Surprisingly, we find that each gene has a unique response to changes in [TF] and promoter activity. As [TF] increases, some genes increase in expression faster than expected from the promoter alone, which we call Promoter Activity Amplification (PAA), while other genes increase in expression more slowly, which we call Promoter Activity Buffering (PAB). We call the general phenomena Promoter Activity Buffering or Amplification (PABA). Sequences outside of the promoter buffer, or amplify, changes in promoter activity. While one possible mechanism for PABA is autoregulation, autoregulation is not responsible for PABA in native genes. Experimental data and mathematical modeling suggests that titration of limiting *trans* regulators is responsible for PABA in some native genes. Finally, we show that, because promoter activity buffering reduces the effect of changes in [TF] on gene expression, buffering insulates cells from the potentially deleterious effects of misregulation.

## RESULTS

### Sensitivity to changes in [TF] is influenced by sequences external to the promoter

To directly test if a gene’s sensitivity to changes in [TF] is influenced by elements external to the promoter **(Figure 1A)** we generated a master strain in which [TF] can be varied without affecting any other cellular processes(McIsaac et al., 2013), and in which promoter activity and protein expression level can be independently measured(Schikora-Tamarit et al., 2016a). The master strain contains the engineered *β*-estradiol activated Z_3_EV transcription factor and GFP driven by a Z_3_EV regulated promoter (Z_3_EVpr). From this master strain we generated a library of 66 strains. In each strain a single gene at its native locus had its native promoter replaced with the Z_3_EV promoter driving the expression of a mCherry-gene construct **(Figure 1B)**. For each gene we used flow cytometry to measure how promoter activity (Z_3_EVpr-GFP) and protein expression level (mCherry) change as function of [TF] (*β*-estradiol) **(Figure 1C)**. Of the 66 tagged genes, 42 gave reproducible results across multiple independent transformants and were used for further study **(Supplementary Table 1)**. We found that, in spite of having identical promoters, different genes exhibit vastly different response to changes in both [TF] and promoter activity **(Figure 1C,D, Figure S1, S2, S3, S4)**.

**Figure 1:**
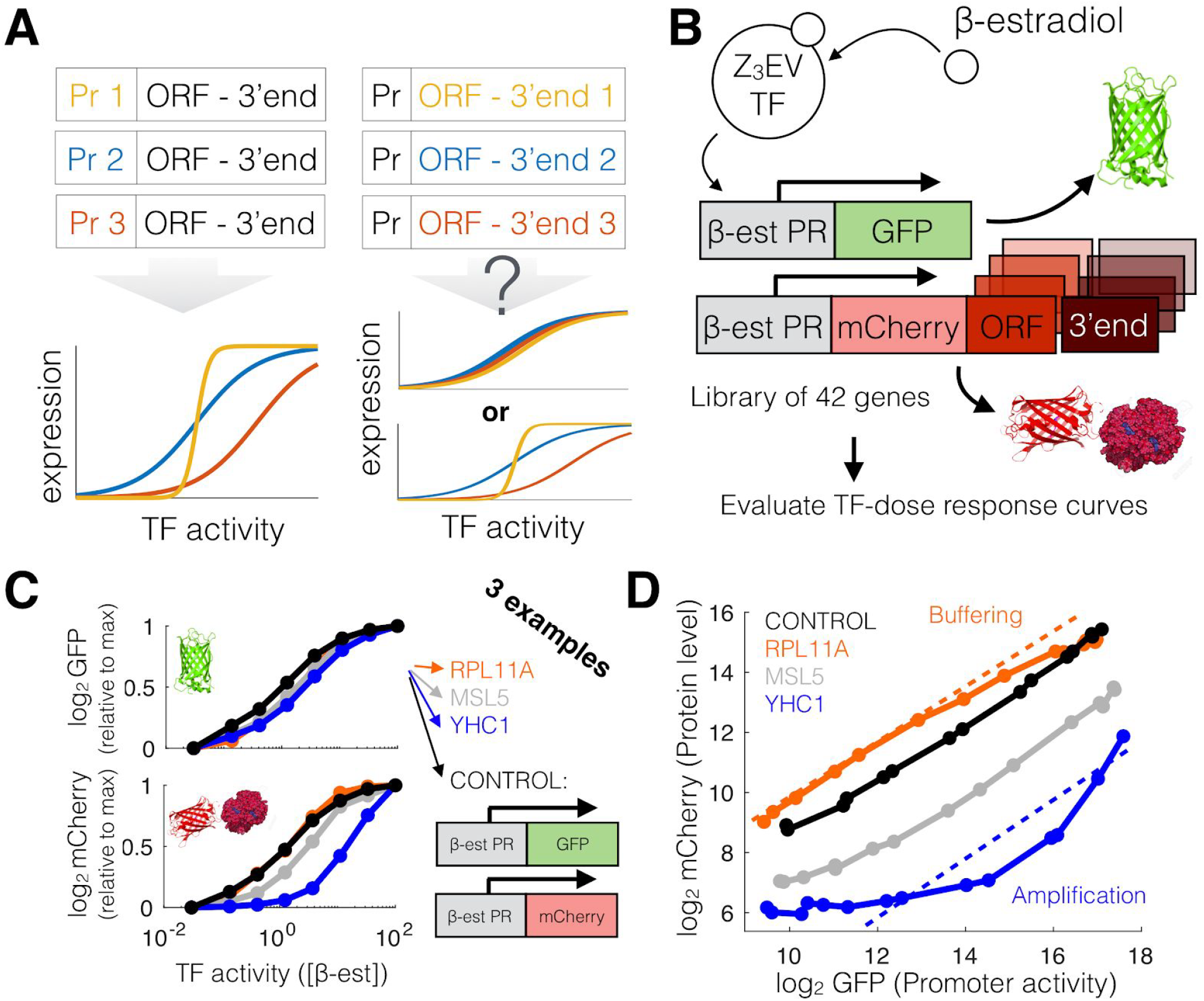
A synthetic gene circuit for determining how TF Dose-Response Curves (TF-DRCs) depend on sequences external to the promoter. **(A)** A schematic view of the question. Promoter elements determine the response to Transcription Factor (TF) activity in gene expression. However, it is unknown which is the contribution of the Open Reading Frame (ORF) and 3’end regions. **(B)** As an example, *β*-estradiol activates the engineered zinc-finger TF Z3 (McIsaac et al., 2013). We engineered a yeast strain library of 42 genes, replacing the native 5’ regulatory region with a synthetic *β*-estradiol promoter driving an mCherry-protein tag, while GFP remains as a pathway-specific reporter of promoter activity of each gene. **(C)** Three example genes and a control strain (that lacks an mCherry-protein tag) show similar TF activity-vs-GFP expression profiles (top). However, TF-vs-mCherry-protein expression profiles (bottom) are diverse. **(D)** Each gene exhibits a unique GFP (as a proxy for promoter activity) vs mCherry (as a proxy for protein level) profile, despite having identical promoters. Three example genes illustrate the diverse shapes of the curves.

In comparison to a control strain in which Z_3_EVpr-mCherry is integrated at the *his3* locus, we found three types of behavior **(Figure 1D)**. For some genes, such as MSL5, expression is offset from the control strain by a constant amount. For other genes, such as RPL11A, at high promoter activity expression increases more slowly than in the control strain. We refer to this as Promoter Activity Buffering. Finally, for some genes, such as YHC1, at high promoter activity, expression increases more rapidly than in the control strain. We refer to this as Promoter Activity Amplification.

### A mathematical model to quantify Promoter Activity Buffering or Amplification (PABA)

To quantify Promoter Activity Buffering or Amplification (PABA) we built a mathematical model that describes how protein levels change with increasing promoter activity **(see Methods)**. The model contains three mCherry-protein-specific biological parameters: (1) protein and mRNA synthesis and degradation rates (SD), (2) basal expression (BE, measured as the fraction of maximum fluorescence in the absence of *β*-estradiol) and (3) expression (protein) level dependent regulation of expression (F, which allows protein levels to feedback on and influence transcription or translation rates) **(see Materials and methods)**. PABA quantifies protein expression level dependent regulation of gene expression. Changing each parameter has a unique effect, allowing us to quantitatively isolate the effect of PABA on gene expression, and to predict what expression would look like in the absence of PABA **(Figure 2A,B)**.

**Figure 2:**
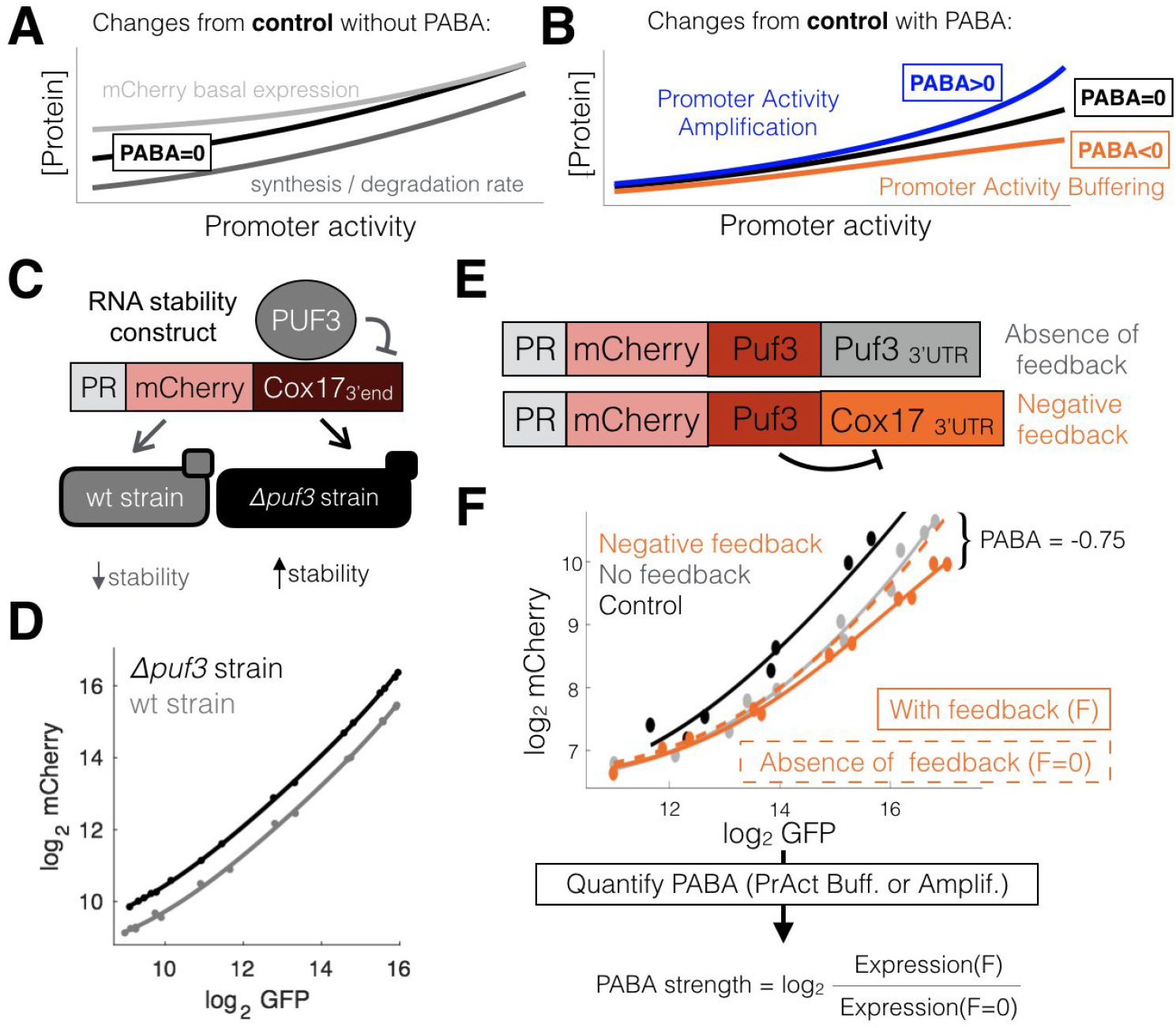
A mathematical model for quantifying Promoter Activity Buffering or Amplification, such as those produced by autoregulatory feedback loops. **(A)** Any gene measured in our library depicts a particular GFP (Promoter activity) vs mCherry ([Protein]) curve. A minimal model of gene expression, together with experimental data (see **Supplementary information)**, predicts that changes in mCherry basal expression and/or synthesis/degradation rates can alter the basal curve (black) in the absence of Promoter Activity Buffering or Amplification (PABA). **(B)** In contrast, autoregulation models Amplification (PABA > 0, blue) or Buffering (PABA < 0). **(C)** As an example of (A) we built the *Z_3_EVpr-GFP, Z_3_EVpr-mCherry-Cox17_3’End_* construct into *wt* an*d puf3 Δ* strains. As Puf3p destabilizes the Cox17_3’end(Olivas and Parker 2000)_, each of the genetic backgrounds should depict a different mCherry RNA stability. **(D)** Shown are measured GFP and mCherry of these constructs at different *β*-estradiol concentrations. **(E)** As a proof of (B) we used the previously described (Schikora-Tamarit et al., 2016b) set of negative feedback strains. These are built in a *Z_3_EVpr-GFP* background and express either a *Z_3_EVpr-mCherry-Puf3-Cox17_3’END_* or a *Z_3_EVpr-mCherry-Puf3-Puf3_3’END_*, resulting in presence and absence of negative feedback. **(F)** Expression data (dots) of these constructs (together with the control strain in black) is fit (full lines) by a model that implements changes in mCherry basal expression and synthesis/degradation rates in the *Puf3-Puf3_3’END_*, but PABA (computed as F (feedback)) is required to fit the *Puf3-Cox17_3’END_* strain. However, setting F to 0 in the *Puf3-Cox17_3’END_* (dashed orange line) models a curve that fits the *Puf3-Puf3_3’END_* data. We thus calculate PABA strength as the mCherry expression ratio of these two models (with (F) and without (F = 0) PABA). It has to be noted that all the Puf3 constructs were engineered using a not yeast codon optimized mCherry, with the corresponding fluorescence control strain.

Negative autoregulation provides a conceptual and experimental model for Promoter Activity Buffering. In the presence of negative autoregulation, increased protein levels reduce transcription or translation rates, resulting in less protein than expected in the absence of autoregulation **(Figure 2B)**. Conversely, positive autoregulation results in a faster than expected increase in protein levels, and hence results in promoter activity amplification (PAA). To quantify PABA and to test the model we took advantage of the RNA binding protein Puf3, which binds to sequences in the COX17 3’UTR, stimulating degradation of the mRNA (Olivas and Parker, 2000). We generated a strain in which the Z_3_EVpr-mCherry construct contains the COX17 3’end (3’UTR + transcription terminator). Compared to a *PUF3^+^* strain, mCherry protein expression in a *puf3 Δ* strain is increased by a constant offset, consistent with the model **(Figure 2A,C,D)**. In contrast, a negative autoregulation strain in which the native *PUF3* gene is replaced by Z3EVpr-mCherry-PUF3-COX17_3’end_ exhibits Promoter Activity Buffering, with respect to a Z3EVpr-mCherry-PUF3-PUF3_3’end_ strain **(Figure 2E,F)**. Thus, constant changes in mRNA stability do not buffer protein levels from changes in promoter activity. In contrast, changes in mRNA stability as a function of promoter activity result in PABA in this well-defined synthetic system.

To quantify PABA for each gene we first fit the model to expression data from the control strain. We next allow either SD and BE to vary or SD, BE, and F to vary. Variation in SD and BE are sufficient to fit the no-feedback strain (R^2^=0.998), but F is required to fit the negative-feedback strain(SD,BE R^2^=0.995; SD,BE,F R^2^=0.999). While this increase in R^2^ is small, it is consistent across biological replicates on a model trained on one biological replicate and used to predict a second biological replicate. We then computationally removed feedback by setting F to zero, and calculate PABA strength as the change in protein expression at maximal induction when F is set to 0. Consistent with our model, computational removal of PABA by setting F to 0 results in an accurate prediction of expression in the no-feedback strain **(Figure 2F)**.

### Promoter Activity Buffering or Amplification is common and continuous

To quantify the prevalence of PABA and to predict what expression would look like in the absence of PABA, we fit the model to Z_3_EVpr-mCherry-expression data for 42 native yeast genes. Our model fits experimental data of all genes **(Figure S5)**. We find that PABA is not binary, but exists in a continuum, from strong buffering to strong amplification, with promoter activity buffering being the most prevalent **(Figure 3)**.The dynamic range of expression observed in our experimental system includes native expression levels **(Figure S6)**, suggesting that PABA acts in intact native regulatory networks. We note that there is no correlation between PABA and expression level **(Figure S7)**, suggesting that PABA is not due to a general toxicity of overexpression. Buffering can result in more than a 75% reduction (−2 fold change) in expression levels, while amplification can result in a more than a 2 fold increase in expression **(Figure 3)**. These changes in expression are well within the range that natural selection can act on gene expression(Wagner, 2005; Zeevi et al., 2014).

**Figure 3:**
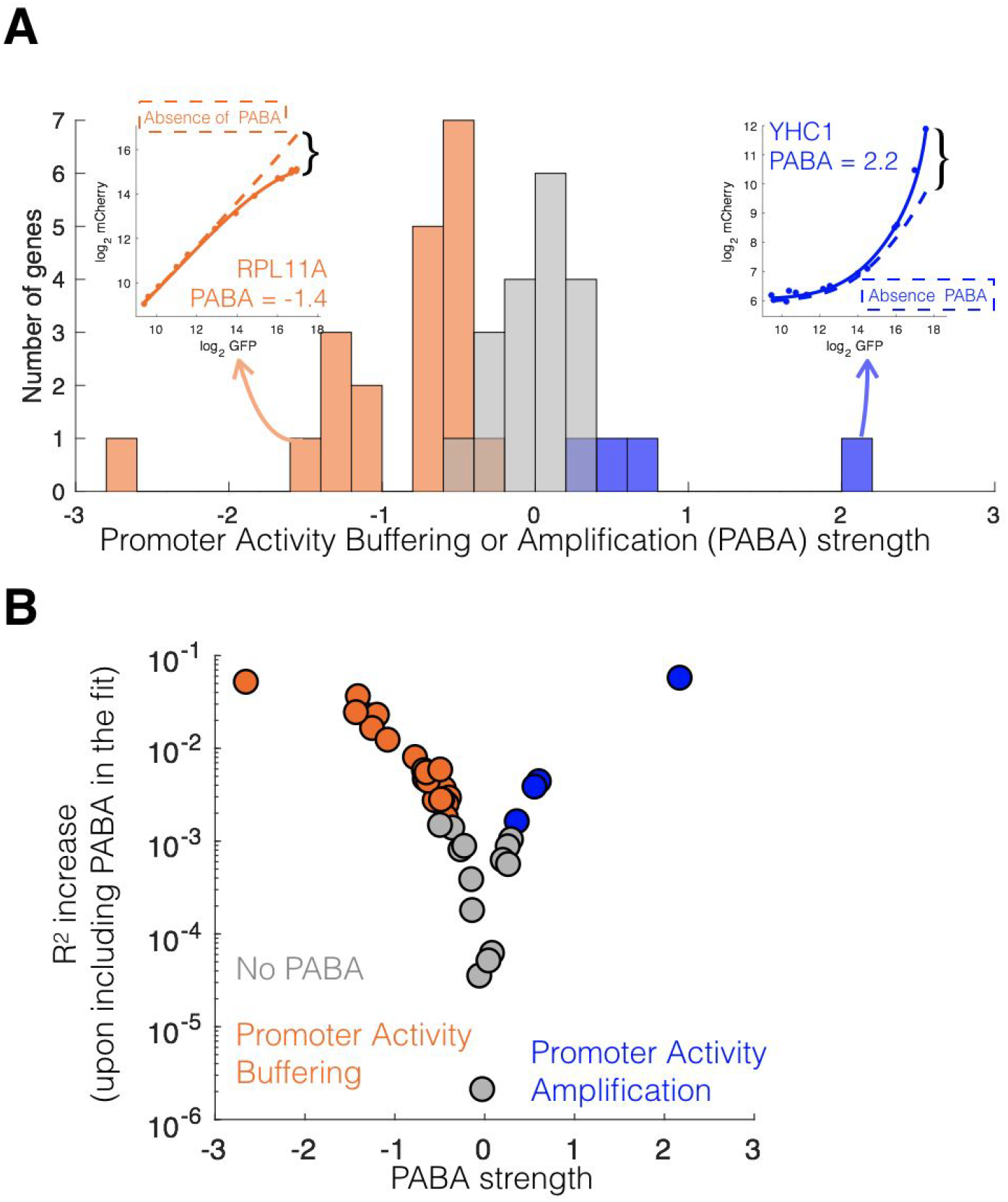
Promoter Activity Buffering or Amplification (PABA) is common and continuous. **(A)** We measured GFP and mCherry expression in our library (small insets represent two example genes) and quantified PABA strength on them as described above (where dashed lines represent the absence of PABA). Genes can be classified into promoter activity buffering (orange), amplification (blue) and no PABA (gray). **(B)** The R^2^ increase of a model that includes PABA with respect to the basal fit (just including non-PABA parameters to explain the data) represents the relevance of the fit. We set a threshold for relevance (see methods) to assign the presence of PABA (orange for Buffering, blue for Amplification and gray for no PABA. This is also related to (A)) for each gene.

The forty-two measured genes cover a diverse class of RNA binding proteins involved in splicing, regulation of both transcription and mRNA turnover and ribosome biogenesis. Twenty-three of our genes have orthologs involved in human disease **(Supplementary Table 1)**. We find that all (5/5) of the genes involved in ribosome biogenesis (Nop12, Nop4, Rpl11A, Rpl11B and Tif4631) have promoter activity buffering (GO term enrichment, Fisher’s exact test, p=0.019), while all genes involved in mRNA processing (Rna15, Smx3 and Yhc1) have promoter activity amplification (p=0.037).

### ORF and 3’end sequences encode Promoter Activity Buffering or Amplification

PABA, by definition, is promoter-independent. In order to give further mechanistic insights into PABA we tested if it is encoded in the open reading frame (ORF) or 3’ end (3’UTRs and transcription terminator) **(Figure 4A,B)**. To do so we generated strains in which the native 3’ end of each gene was replaced with the *Ashbya gossypii* TEF 3’end, resulting in a set of *gene::Z_3_EVpr-mCherry-gene_ORF_-TEF_3’END_* constructs. The genes chosen included strong promoter activity buffering (Rpl11A, Hek2, Nhp2, Rrp40) strong amplification (Yhc1) or no PABA (Puf2). We compared the expression of these strains with the corresponding *gene::Z_3_EVpr-mCherry-gene_ORF_-gene_3’END_* strain. Any change in PABA implicates the 3’end. Changing the native 3’end modifies induction curves in a gene-specific manner **(Figure 4C, S7)**. 4/5 tested genes (Rpl11A, Hek2, Nhp2, Rrp40) lose PABA when their 3’end is replaced, indicating that both the 3’end and the ORF can encode PABA **(Figure 4D)**. In addition, fusion of 3’ends to mCherry confers PABA independently of the location in which the construct is integrated into the genome, though the PABA conferred by the 3’end is not always the same as that conferred by the ORF + 3’end, suggesting that, for some genes, PABA is determined by an interaction between these two features **(Figure S16)**. Importantly, changing the 3’end affects PABA independent of its effect on protein abundance **(Figure 4C, S7)**. Thus, both the 3’ ends and ORFs play a role not only in determining absolute protein expression levels, but also influence the effect of [TF] and promoter activity on gene expression.

**Figure 4.**
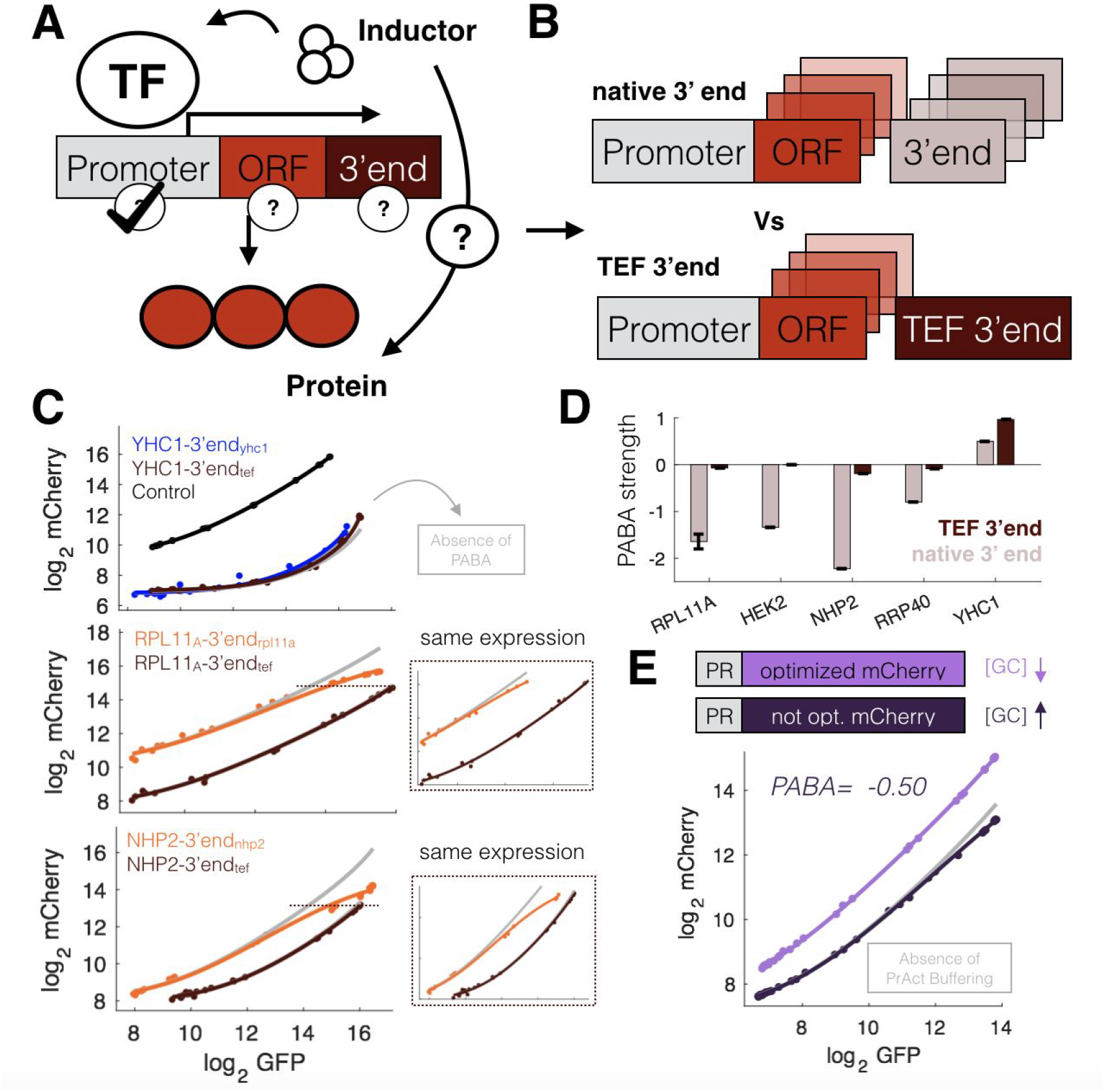
ORF and 3’end sequences encode Promoter Activity Buffering or Amplification (PABA). **(A)** The promoter encodes the TF-response (Carey et al., 2013), but either open reading frames (ORF) and/or 3’end regulatory may also play a role through PABA. **(B)** To test this we compared expression in the library with a set of strains in which the native 3’end was replaced with the *TEF*_*3’EN*D_. **(C)** Three example genes with either Amplification (YHC1) or buffering (RPL11A ans NHP2) preserve or loose the PABA-like shape when the native 3’end is replaced (brown lines). The gray lines represent the absence of PABA for each curve. The small plots correspond to the same analysis performed on equal maximum mCherry expression of the native and TEF 3’end, for RPL11A and NHP2. **(D)** Mean and SEM PABA values obtained in 30 runs of the model. Each assayed gene has a different PABA strength in the native (brown) or *tef* 3’end (light brown) constructs. **(E)** Two fluorescent control strains (with the *β*-estradiol promoter driving GFP and mCherry alone) that differ only by mCherry ORF sequence exhibit different PABA behaviors. An mCherry which is not codon optimized and GC rich depicts Promoter Activity Buffering.

3’ end regions are well-established regulatory regions, but how often coding sequences play an active role in regulating gene expression is less clear(Plotkin and Kudla, 2011). To confirm that the ORF sequence itself can be responsible for PABA we compared the promoter-vs-expression profiles of two mCherry constructs that differ only by synonymous mutations. An mCherry with higher GC content and lower codon bias exhibits promoter activity buffering **(Figure 4E)**, confirming that PABA can be encoded in the ORF.

### Promoter Activity Buffering or Amplification is not mediated through autoregulation

Autoregulation is one mechanism by which PABA could be implemented in native genes. In both native and synthetic regulatory networks (Becskei and Serrano, 2000; Eriksson et al., 2012; Springer et al., 2010), negative autoregulation results in dosage compensation, in which deletion of one copy of a gene results in increased expression of the other copy. Therefore, one way to test for autoregulation is to measure how expression of one allele changes when the other allele in a diploid is deleted. Genes that exhibit promoter activity buffering (PAB) should exhibit dosage compensation if PAB is due to negative autoregulation. The same reasoning can be applied to promoter activity amplification (PAA). To test this we generated diploids in which the other allele of the inducible tagged gene is either wild-type or deleted, and measured protein levels **(Figure 5A)**. Negative autoregulation results in dosage compensation (increased expression) when the other allele is deleted **(Figure 5B)**. The effect is especially strong at low promoter activity, where the native allele contributes more to the total cellular protein concentration. However, no native genes exhibited gene copy-number dependent effects on expression **(Figure 5C)**, suggesting that PABA is independent of autoregulation.

**Figure 5.**
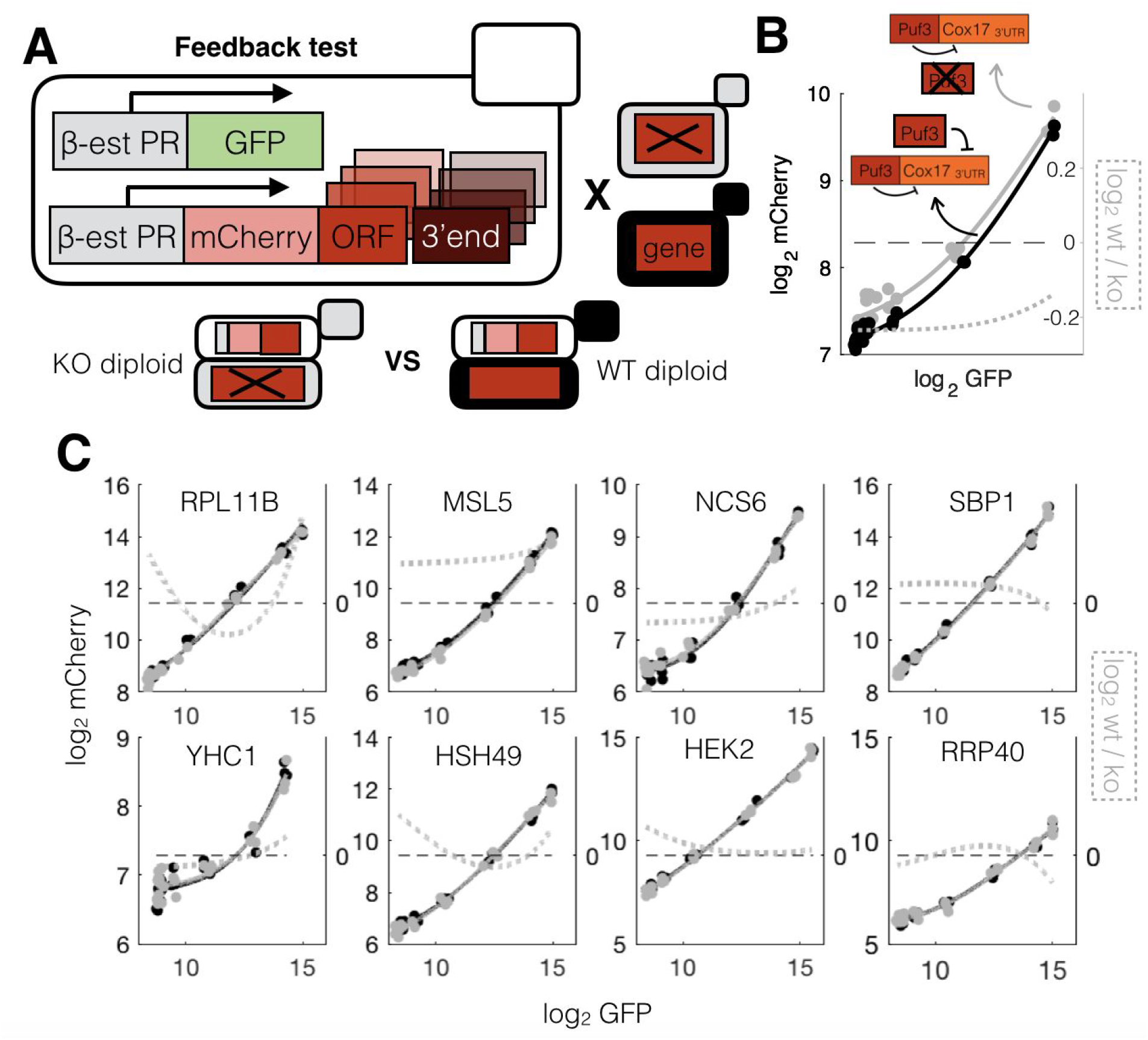
Promoter Activity Buffering or Amplification (PABA) does not necessitate feedback regulation. **(A)** We generated strains to test if PABA requires feedback regulation mating the library strains with either a *Δgene* (gray) or a *wt* (black) strain. The resulting diploids are hereafter referred as KO or WT strains. The idea is that if feedback exists, adding an extra copy of the gene must have an effect on mCherry expression. **(B)** The *puf3-cox17_3’END_* construct has higher mCherry expression in the KO diploid, as the higher Puf3 abundance in the WT diploid results in less mCherry-Puf3. The dashed line represents mCherry expression ratio between the two strains. **(C)** In contrast, none the strains engineered genes (bearing high buffering, amplification or no PABA in the haploid library) depicts any visual difference in mCherry expression between the two diploids.

### Titration of limiting regulators can explain PABA in synthetic constructs

The observation that PABA is mostly encoded in the 3’end of genes and that negative autoregulation acting on mRNA stability results in PABA suggested that mRNA stability might be responsible for PABA in native genes. To test this we searched a dataset of 90 gene-specific features such as codon bias, amino-acid properties and 3’UTR length for features that correlate with PABA. We found that mRNA stability-related properties (including codon bias and UTR features) are the only predictors of PABA strength. **(Figure S15)**. As the correlation of PABA and mRNA stability is negative **(Figure S15)**, Promoter Activity Buffering may be a property of stable transcripts, while Amplification may be a property of unstable transcripts. Cell-to-cell variability data for some genes are consistent with a change in mRNA stability **(Figure S15)**, though steady-state protein levels are not sufficient to uniquely identify changes in mRNA stability (Baudrimont et al., 2017; Thattai, 2016; Zenklusen et al., 2008). It is likely that many other biological processes, such as transcription elongation, transcription termination or protein-complex imbalance-mediated protein degradation can also play a role, and the precise biological factors that determine PABA will be unique to each gene.

The fact that PABA can be encoded in either the open reading frame or the 3’end **(Figure 4)** suggests that, for each gene, gene-specific mechanisms are involved. We therefore hypothesized one instantiation of a model for PABA. In this model translation changes as a function of promoter activity due to gene-specific *trans* regulators being present in limiting amounts. If a positive regulator becomes limiting at high promoter activity, the rate of increase in protein expression will decrease, resulting in promoter activity buffering. The opposite trend would be observed in a transcript that is unstable due to a limiting *trans* regulatory factor. We call this the “*trans*-element titration” model for PABA. **(Figure 6A)**.

**Figure 6.**
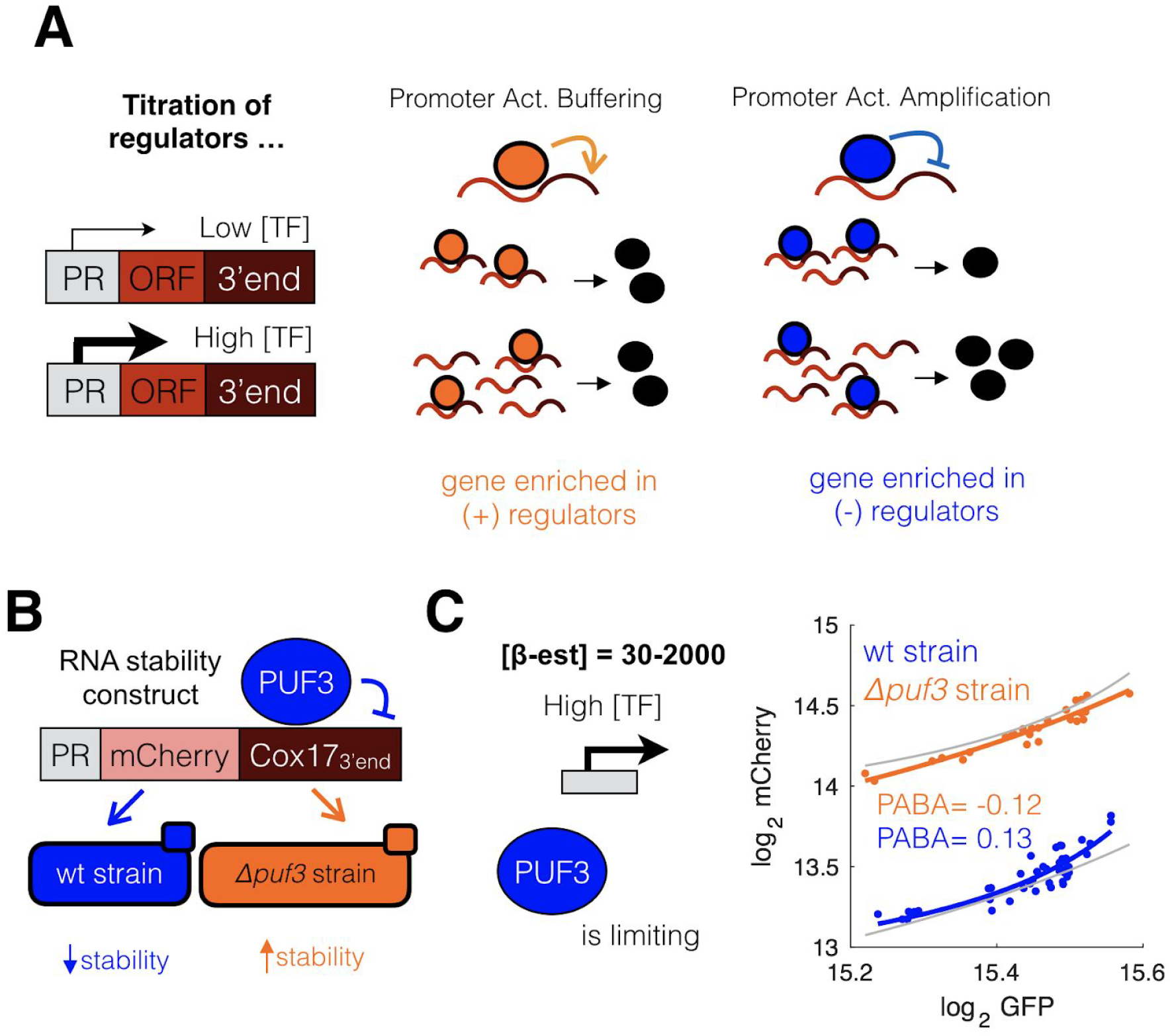
Titration of limiting regulators explains Promoter Activity Buffering or Amplification (PABA). **(A)** A possible mechanism for PAB is that genes enriched in positive regulators (of transcription, mRNA degradation and/or translation) are efficiently translated at low induction levels. However, when the induction level is high, the loading becomes limiting and some mRNAs undergo inefficient transcription, degradation and/or inefficient translation because of titration of the regulator to other transcripts. The opposite would be observed in PAA, as there’s an enrichment in negative regulators. **(B)** A *cox17::Z_3_EVpr-mCherry-cox17_3’END_* depicts higher mRNA stability in a *Δpuf3* (orange) compared to a *wt* (blue) strain (see **Figure 2C,D**), as Puf3 is a destabilizing regulator of the *cox17* 3’UTR (Schikora-Tamarit et al., 2016a). **(C)** At high induction mCherry-COX17_3’end_ exhibits PAA in a PUF3^+^ strain but not in a *puf3 Δ* strain, suggesting that Puf3p may be limiting. The gray lines show the offset curve of the other strain.

To test this idea we used a *cox17::Z_3_EVpr-mCherry-COX17_3’END_* strain in which Puf3 is limiting. At high promoter activity the mCherry mRNA should become increasingly stabilized due to Puf3 titration, resulting in promoter activity amplification. Experimental data **(Figure 6B,C, Figure S14, S15)** are consistent with the titration model.

### Titration of limiting positive regulators is a potential mechanism for ploidy-dependent dosage compensation

We found that changing gene copy number in a diploid does not change expression of any of the *Z_3_EVpr-mCherry-gene* constructs **(Figure 5)**, which might reveal an interesting behavior in the context of a limiting-regulator titration model. We modelled **(see Methods)** the induction profiles of *Z_3_EVpr-mCherry-gene* constructs by changing the sign of regulation (positive or negative) and the concentration of regulator ([R]), which results in PABA. Surprisingly, our model predicts that doubling the amount of [R], as would happen in a hemizygous diploid with only a single copy of the inducible gene, results in a reduction of PABA towards zero **(Figure 7A)**.These findings suggest that PABA (which corresponds, by definition, to dosage-sensitive changes in expression) depends on ploidy in the context of a limiting-factor titration.

**Figure 7.**
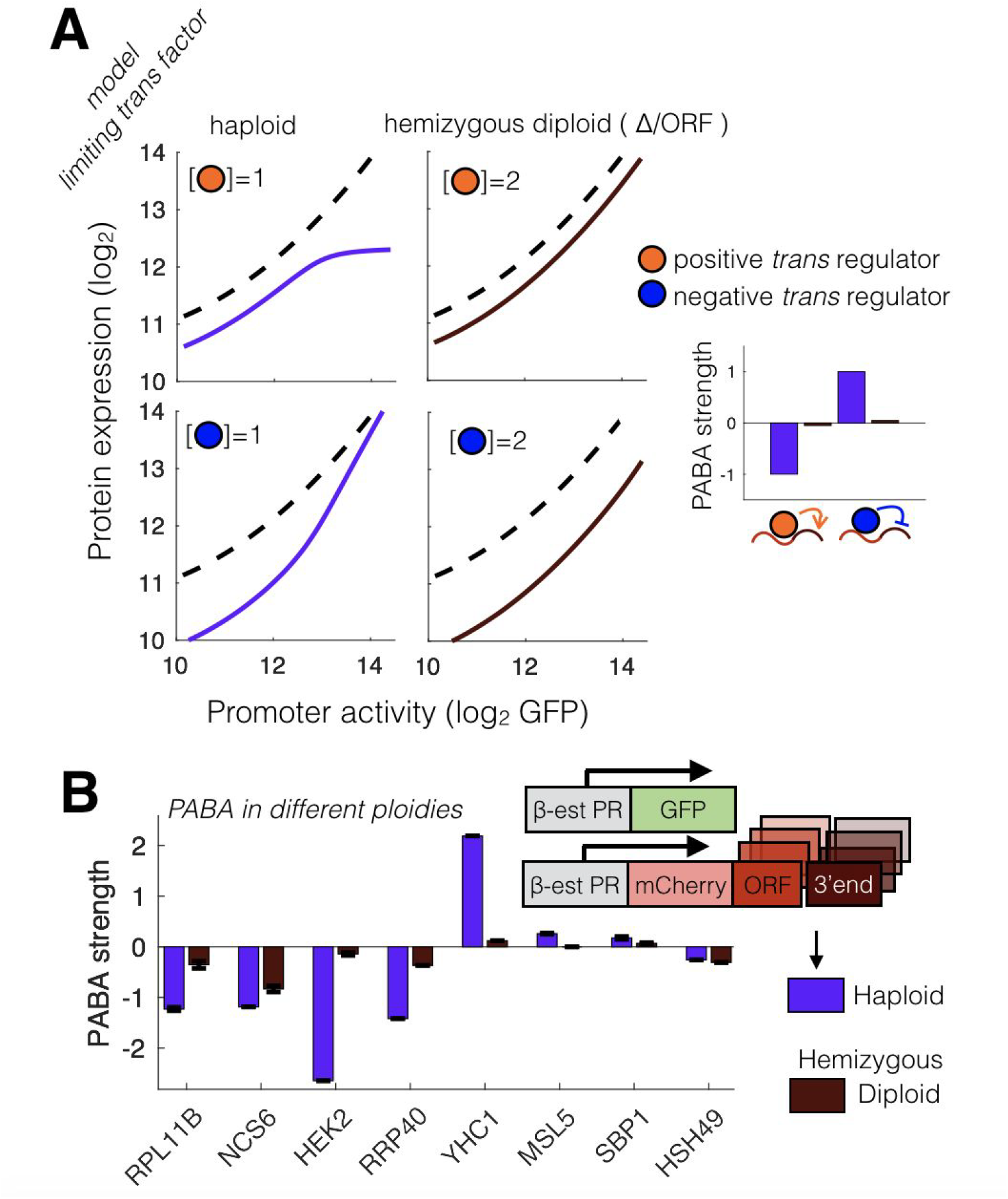
Titration of limiting regulators is a potential mechanism for ploidy-dependent dosage compensation. **(A)** In a model of gene expression that implements PABA through titration of limiting regulators, the mCherry vs GFP profile of any gene in our library should be affected by both sign of the regulation and ploidy. The upper panels show these profiles when a positive regulator generates Promoter Activity Buffering, and vice-versa (a negative regulator generates Promoter Activity Amplification) in the bottom panels. The right panels show the model in a hemizygous diploid, where the amount of regulator is doubled. The right-most plot shows the PABA strength of each of the four simulations. **(B)** Measured PABA strength in the haploid and hemizygous diploid strains. Error-bars refer to standard error of the mean of the PABA values obtained in 30 runs of the model fit to experimental data.

To test this prediction we generated diploid strains expressing the *Z_3_EVpr-mCherry-gene* constructs. Consistent with the model, PABA is reduced in diploids, and this reduction is stronger for native transcripts, in which PABA is likely due to titration of limiting *trans* regulator (s), than for the Puf3-COX17 negative autoregulation control strain. **(Figure 7B)**. As Promoter Activity Buffering (PAB) is the most common phenomena, we conclude that titration of limiting positive regulators is a potential mechanism for ploidy-dependent dosage compensation.

### PABA modifies the fitness cost of promoter activity

Protein levels of many genes are tightly regulated, and changes in expression level can lead to changes in fitness (Keren et al., 2016; Rest et al., 2013). Our data suggest that the relationship between promoter activity and expression is gene-specific, and therefore protein expression levels cannot be precisely inferred from measured promoter activity. We therefore sought to determine the mapping between fitness and actual protein levels. A simple model in which expression affects fitness suggests that PABA should modify the fitness effects of changes in [TF] and that promoter activity buffering also buffers the fitness cost of overexpression **(Figure 8A)**.

**Figure 8.**
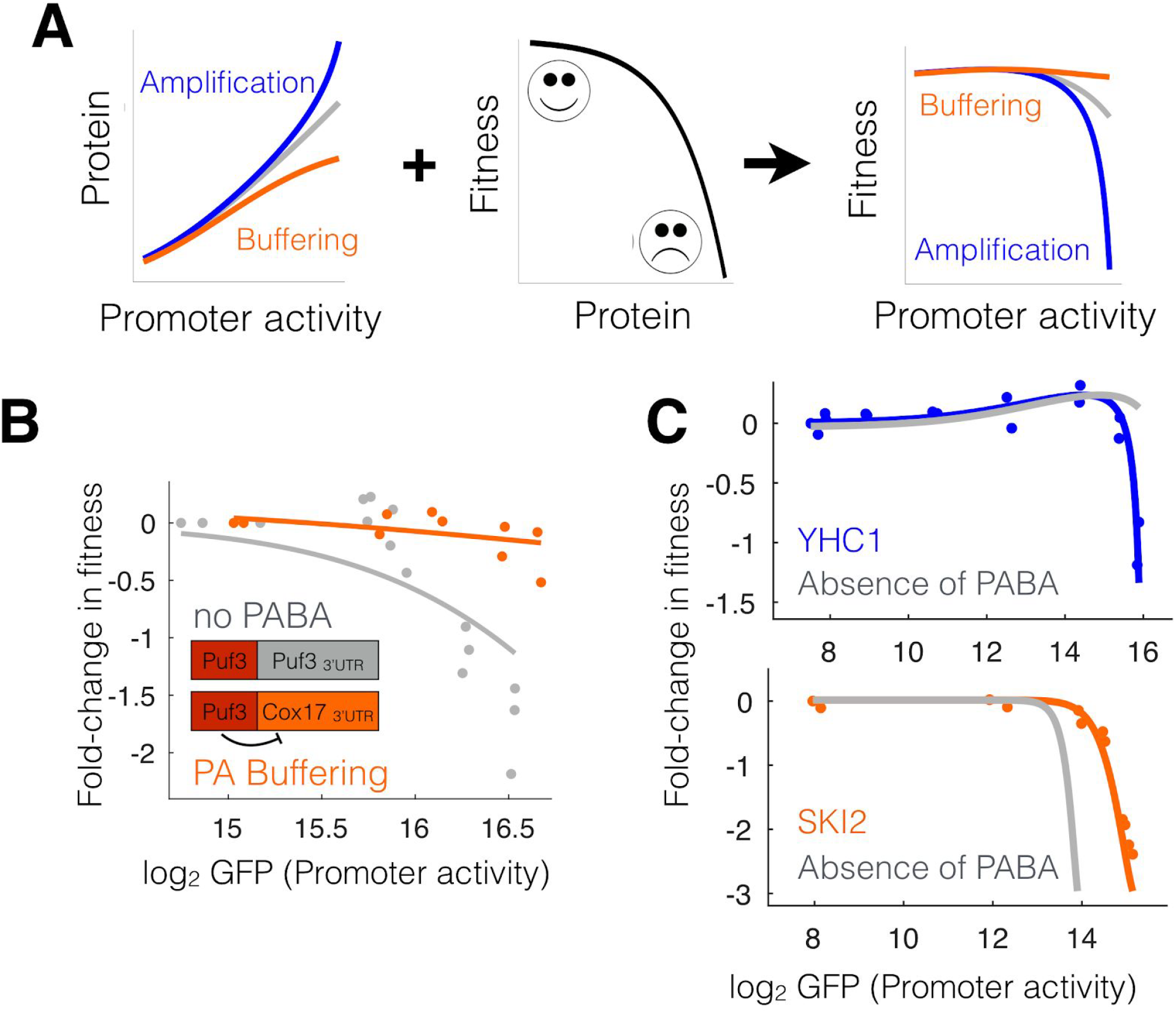
PABA modifies the fitness cost of excessive promoter activity. **(A)** Genes are subjected to Promoter Activity Buffering or Amplification (left), and overexpression of many proteins can be deleterious for the cell (middle, (Keren et al., **2016))**. According to a simple model that relates promoter activity to protein expression and fitness, PABA can enhance or buffer cells from the fitness cost of excessive promoter activity. **(B)** As a proof of concept, we measured the Fold-change in fitness of the two *Z_3_EVpr-mCherry-Puf3-Cox17_3’END_* and *Z_3_EVpr-mCherry-Puf3-Puf3_3’END_* constructs, at different induction rates. These correspond to the presence and absence to Promoter Activity Buffering, respectively. Experimental data (dots) is well fit by an impulse function model (lines) as previously described (Chechik et al., 2008). **(C)** Shown are two example genes that have a fitness defect at high expression (dots are experimental data and colored lines are the model fit) have a different relationship between promoter activity and fitness in the absence of PABA (gray lines, see **Methods**).

To test this hypothesis we measured fitness at different induction levels of the Puf3-COX17_3’end_ synthetic negative autoregulation strain using a flow-cytometry based competition assay **(see methods)**. We found that Puf3 lacking negative autoregulation (and lacking PAB) exhibits a large fitness defect at high promoter activity levels. In contrast, the strain with negative autoregulation exhibits promoter activity buffering and no fitness defect. **(Figure 8B)**.

To determine the relationship between fitness and protein expression in native genes we measured fitness for 12 strains **(see methods)**. Five genes exhibited a large decrease in fitness at high expression, one exhibited an increase in fitness at high expression, and three exhibited no fitness change **(Figure S8)**. Interestingly, the two 60S ribosomal paralogous proteins, RPL11A and RPL11B, (homologous to the mammalian L11 and bacterial L5 proteins) both exhibit a non-monotonic relationship between expression and fitness **(Figure S8)**, as well as promoter activity buffering, suggesting that tight regulation of expression of these genes is essential, and that this tight regulation is mediated by PABA.

To determine if PABA alters the mapping between promoter activity and fitness we used our mathematical model to predict protein expression in the absence of PABA for native genes. We find that, if a gene has a fitness defect due to overexpression, promoter activity buffering can increase up to ~2 fold the induction level at which the cells exhibit maximal fitness. On the other hand, promoter activity amplification results in a large growth defect **(Figure 8C)**. Therefore, by modifying the mapping between promoter activity and protein expression, PABA changes the fitness landscape of gene expression.

## DISCUSSION

By placing diverse native genes at their native genomic locus all under the control of a single inducible promoter and measuring how protein levels changes as a function of promoter activity, we found that the shape of a gene’s TF dose response curve is affected by sequences outside of the promoter. The result is that many genes exhibit promoter activity buffering, in which the protein abundance increases more slowly than would be expected from the amount of active transcription factor inside the cell. This effect is not due simply to differences in mRNA or protein stability between genes. Instead it is likely because the molecular mechanisms that regulate gene expression change throughout the induction curve. This effect is often encoded in the 3’end of the gene, but can be encoded in the open reading frame as well. While this effect can be implemented by autoregulatory feedback loops, this does not seem to be common in native genes. Instead it is likely that titration of limiting *trans* factors plays a major role.

Our data suggest that protein expression levels for more than half of genes are influenced not only by transcription factor concentration and activity, but also by the copy number of both the gene of interest, and the copy number and expression level of other genes in the cell (Brewster et al., 2014; Buchler and Louis, 2008; van Dijk et al., 2017). This includes both *trans* regulators of transcription and translation, other targets of these *trans* factors, and the copy numbers of physical interaction partners of the protein (Gonçalves et al., 2017; Ishikawa et al., 2017; Kim and Hart, 2017; Ryan et al., 2017; Sung et al., 2016). One possible mechanism that can explain ORF-encoded PAB, but not obviously PAA, is that proteins that are overexpressed compared with other members of a complex are degraded (McShane et al., 2016). When this is due to variation in gene copy number, as in the case of a hemizygous diploid, PAB can result in dosage compensation, rescuing haploinsufficiency and providing genetic robustness.

PAB in the mCherry with high GC content and poor codon usage could be explained by the titration of limiting factors models. From the perspective of codon usage, this transcript looks nothing like highly expressed genes in yeast (Harigaya and Parker, 2016; Radhakrishnan et al., 2016). It is possible that some critical factor, such as a rare tRNA, is titrated away at high expression. Transcriptional elongation may also play a role; the high GC mCherry is predicted to have higher nucleosome occupancy (Kaplan et al., 2009). This suggests the existence of a general mechanism to dampen expression, possibly by slowing transcription or translation, and/or accelerating mRNA degradation, when a sub-optimal gene is overexpressed.

The need for PAA, which is less common, is less obvious, as amplification decreases robustness **(Figure 8)**,. We note that large Hill coefficients are not uncommon in biology, and that both positive feedback loops and coherent feed-forward loops provide strong non-linear responses to changes in an input signal (Shoval and Alon, 2010).

We note that while the magnitude of the effects are often small, PABA does measurably reduce the fitness cost of misexpression; Z_3_EVpr-Puf3 strains lacking PAB have a fitness defect when promoter activity is high. Furthermore, it is important to remember that natural selection will act on mutations that affect fitness (selection coefficient) by more than 1/(2 * effective population size) (~ 10^7^ in yeast) (Kimura, 1983; Tsai et al., 2008; Wagner, 2005). This results in very strong evolutionary conservation of gene expression across species. The median difference in expression level between *S. cerevisiae & S. mitake* ribosomal promoters is less than 10% (Zeevi et al., 2014). In the lab we can, at best, measure fitness effects of 0.1%. It is likely that immeasurably small changes are the dominant force shaping evolution in microbes.

Protein overexpression and underexpression are often deleterious (Keren et al., 2013; Kintaka et al., 2016; Mnaimneh et al., 2004; Rest et al., 2013; Sopko et al., 2006), and yet gene expression and [TF] are noisy (Carey et al., 2013; López-Maury et al., 2008; Nevozhay et al., 2009). Promoter activity buffering makes cells more robust to short-term fluctuations in [TF] as well as more long-term variation in gene copy number. Regulation of protein expression as a function of promoter activity, but not constant changes in mRNA or protein stability, allows genes to be both highly sensitive to regulated variation in [TF] without incurring a fitness penalty from having too little or too much protein expression.

We identified PABA using a theory-first approach (Phillips, 2015). In a mathematical model of gene expression in which changes in [TF] regulate expression, and genes differ from each other only by translation efficiency, mRNA or protein stability, all induction curves will have the same shape. Experimentally, we found that this is not true. Thus identification of PABA shows the strength of theory-inspired experimentation. Likewise, both the hemizygous diploid experiments **(Figure 5)** haploid vs diploid experiments **(Figure 7)** were performed to test predictions made by the autoregulation and titration models, respectively. The results in **Figure 7B** are in line with a rather surprising prediction of the model, and we would not have thought to do the experiment without the model prediction **(Figure 7A)**. There are clearly many unknowns, such as the precise molecular mechanism(s) responsible for PABA at each gene. It is the area in which experiment and model disagree that leaves the most to be discovered.

## Author Contributions

L.B.C. and M.A.S.T. conceived the project. M.A.S.T, C.G.N., G.L.G.S., I.C., X.M.F and M.S. performed experiments. M.A.S.T. developed the models and analyzed the data. L.B.C. and M.A.S.T. wrote the manuscript with input from all authors.

## Acknowledgments

L.B.C. was supported by Ministerio de Economía y Competitividad (MINECO) (BFU2015-68351-P) and AGAUR (2014SGR0974 & 2017SGR1054), and the Unidad de Excelencia María de Maeztu, funded by the MINECO (MDM-2014-0370). We’d like to thank the UPF/CRG Flow Cytometry core. We’d like to thank Alsu Misarova, Carlos Toscano, Abhyudai Singh, Wenfeng Qian, Diego Oyarzun and Javier Macia for helpful comments.

## MATERIALS AND METHODS

### Yeast strains and media

All yeast strains, plasmids and primers used are listed in **Supplementary Tables 2-4**. As a nonfluorescent control strain we used FY4 (LBCY18), a prototrophic variant of S288C (Brachmann et al., 1998). The parental strain for all *promoter::Z_3_EVpr-mCherrγ-orf-3’end* strains (library strains) is DBY19054 (McIsaac et al., 2013) (LBCY105). To generate the control LBCY212 and all library strains (LBCY331 - LBCY453), we generated a PCR amplicon containing KanMX-Z_3_EVpr-mCherry using LBCPlasmid85 and either primers 1 & 2 (control strain) or 3-86 (for each library strain, respectively), and transformed this amplicon into DBY19054. Compared with our previous experimental system(Schikora-Tamarit et al., 2016b), the yeast codon-optimized mCherry induction profile is more similar to the induction profile of GFP, both at the population level and at the single-cell level **(Figure S9)**. The competitor strain for the fitness measurements (LBCY285) is either a BFP+ GFP+ mCherry-haploid spore (obtained from the cross of LBCY88 and LBCY9) or a *fba1::FBA1-mCherry* strain (LBCY310, resulting from the amplicon of PSR101 with primers 131 and 132 into BY4741). The feedback test *wt* diploids (LBCY478 - LBCY485) were made mating the corresponding haploids with BY4742, and the *knock-out* diploids (LBCY514 - LBCY521) were generated inserting a *tef_pr_-hph-tef_ter_* (hygromycin resistance) cassette from PAG34 with primers 133-140 as reverse (and the corresponding forward primers between 3-86) into the corresponding locus of the *wt* diploids. The 3’end replacement strains (LBCY553 - LBCY558) were generated inserting the *tef_pr_-hph-tef_ter_* from PAG34, in the reverse orientation, into the corresponding 3’end region with primers 141-152. As the transcription terminators are almost always bidirectional(Uwimana et al., 2017), the resulting constructs have a constant 3’ end. To generate the *Z_3_EVpr-mCherry-Cox17_3’END_* strains we transformed a KanMX-*Z_3_EVpr*-mCherry amplicon (from LBCP85 and primers 153 & 154) 5’ of the Cox17 3’end in either DBY19054 (*wt*) or LBCY202 (*Δpuf3*). Colony PCR was used to confirm correct integrations of all strains, and to verify that the Z_3_EV promoter has the the correct number of Z_3_EV binding sites. Three - five transformants from each strain were assayed by flow-cytometry to check consistent fluorescent protein expression across transformants to discard the presence of mutations in integrated mCherry PCR amplicon or elsewhere in the genome of the the transformed clone. All transformations were performed using the standard lithium acetate method (Gietz and Woods, 2006). PCR for transformation was performed with Q5 DNA Polymerase (NEB). Colony PCR was performed using Taq Polymerase 2x Master Mix (Sigma Aldrich). Selection for drug resistant transformants was done on YPD plates with CloNAT (Werner bioreagents), G418(VWR) or Hygromycin B (Formedium).

### Flow Cytometry

For all expression measurements, single colonies were picked from YPD plates and inoculated into SCD+0.1 nM *β*-estradiol and grown overnight. Cells were then diluted 1:50 into fresh SCD+0.1 nM *β*-estradiol for a second overnight. Each strain was then inoculated at OD_600_ = 0.001 into different concentrations of SCD + *β*-estradiol (Sigma E8875) and grown for six hours. Strains were then inoculated into fresh SCD + *β*-estradiol at OD_600_ = 0.0012 and grown for an additional 13 hours, (19 hours total growth at 30 ^º^C) and measured by flow cytometry when the OD_600_ ranged between 0.1 and 0.8. Wells in which the cell density was too high or too low were discarded computationally. All flow cytometry was performed on a BD LSRFortessa (BD Biosciences) with 488 nm, 561 nm or 405nm lasers with 530/28, 610/20 or 450/50 filters for GFP, mCherry or BFP, respectively. Analysis of flow-cytometry data was performed using a previously described custom MATLAB pipeline (Carey et al., 2013). To vary the growth rate by varying the carbon source, synthetic minimal media without carbon (SC) was prepared, and then filter-sterilized 20% stock concentration carbon sources (dextrose, sucrose, mannose) were added to the SC.

### Fitness competitions

Competitions in which each inducible RBP strain was competed against the same BFP^+^ GFP^+^ mCherry^−^ or mCherry^+^ competitor strain were performed as follows. Both competitor and RBP were grown for two overnights as above. Then equal amounts of competitor strain and RBP strain were mixed and inoculated, keeping an aliquot of this initial mix for FACS measurement. Fitness at each *β*-estradiol concentration was calculated as the fraction of library strain (calculated as 1 - fraction of cells within an expression distribution corresponding to the competitor strain) after 19 h of growth divided by the same at the inoculation timepoint. Fold-change in fitness (FCF) from the initial induction point was calculated as the log_2_ ratio between fitness at a given *β*-estradiol concentration and fitness at the minimum induction level.

### Mathematical modeling of feedback and fitting to expression data

Based on a simple mathematical model of gene expression, we would expect any protein with a *β*-estradiol induced transcription to have an [mRNA] dynamic as described below:

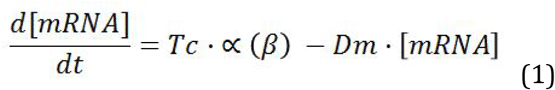

In which Tc is the transcription rate, Dm is the degradation rate of the same mRNA and *α*(*β*) a Hill transfer function that describes the rate of activation of the promoter in response to *β*-estradiol:

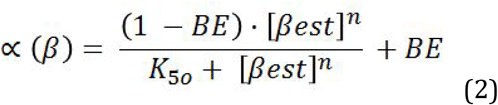

In which n and K_50_ are the Hill coefficient and the [*β*-estradiol] to reach half of the maximum expression, respectively, and BE is the basal rate of activation of the promoter (fraction of maximum expression at [*β*-estradiol] = 0). In terms of protein dynamics, we would expect a behavior such as:

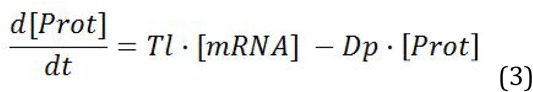

In which Tl is the translation rate and Dp is the protein degradation rate. At steady state (equations 1 and 3 = 0), we would have:

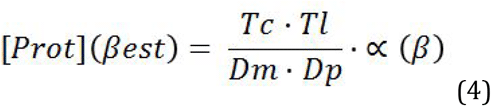

GFP (subscript G) is the control protein for growth, cell size, metabolic and transcriptomic state in our constructs, so we assumed it to follow an induction behavior described by (4), so that:

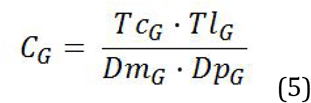

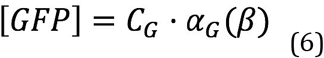

In order to model feedback regulation at the level of transcription or translation we can rewrite of equations (1) and (3) as:

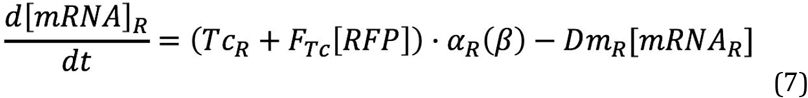

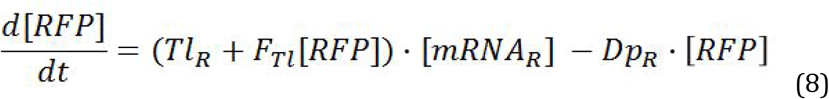

In subscript R refers to mCherry. If there is a feedback effect at the level of synthesis, so that F_Tc_ would be a transcriptional feedback rate, whereas F_Tl_ is a translational feedback rate. On the other hand, a feedback interaction through the degradation would make (1) and (3):

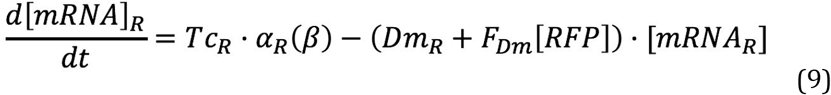

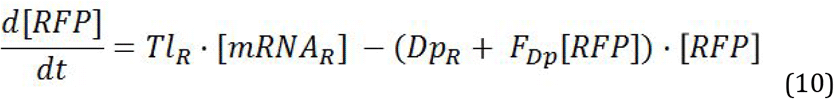

In which F_Dm_ and F_Dp_ are RNA and protein degradation-associated feedback rates, respectively. Simulating these four feedback-model (equations (7) - (10)) at steady state showed that all 4 types of interactions would lead to a similar RFP induction curve at different F values **(Figure S10).**, We therefore only considered a model in which protein synthesis is feedback regulated (equation (8)). Accordingly, for equation (8) at steady state, we have:

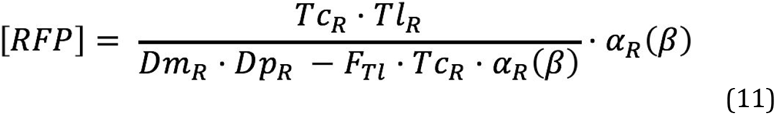

Fitting equation (4) to Y212 data (control strain with no feedback, so that both GFP and mCherry are supposed to follow such a behavior) we experimentally found the following relationships:

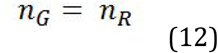

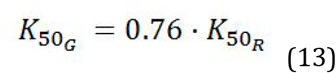

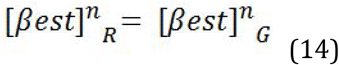

Thus, model which predicts RFP from GFP (RfG) indicates that:

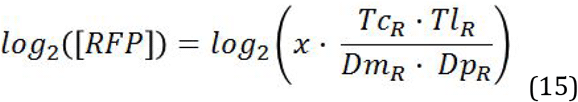

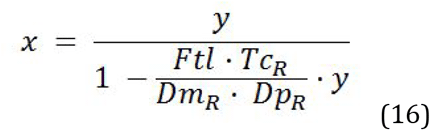

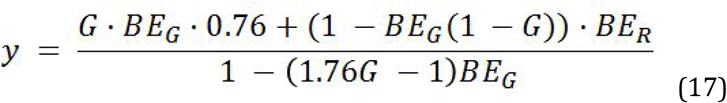

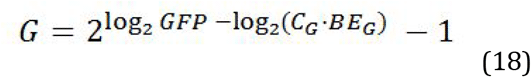

In order to measure feedback strength (or PABA strength) we fit the RfG equations to Y212 mCherry and GFP fluorescence data, setting F_tl_ to 0 and then allowed either BE_R_ and Tl_R_ (as an indicator of synthesis/degradation rates (SD)) or BE_R_, Tl_R_ and F_Tl_ to vary when fitting RBP strain data. These fits correspond to the “basal” and “feedback” parameters, respectively. The absence of feedback was modelled setting F to zero from the “Feedback fit”. We calculate feedback strength (or PABA strength) as the log_2_ ratio of mCherry-protein expression between the presence and the absence of feedback, at maximum induction (100 nM *β*-estradiol). We also computed the difference in R^2^ between the two fits as the relevance of the the feedback measured.

This means that if just allowing BE_R_ and Tl_R_ to vary can explain the data of a construct as well as the case in which F_Tl_ is also changing we say there is no measureable feedback (F is not necessary to explain our experimental data). Note that the GFP parameters were kept constant as a way to set GFP to be the control protein of the system’s activation. Furthermore, we took Tl_R_ as a proxy for SD, as as the only parameter which can be uniquely fit is the Tc*Tl / Dp*Dm ratio **(Figure S10, see Supplementary information)**.

BE_R_ was calculated from a fit (so handled as free parameter) to data and not directly from fluorescence values as the latter method could give a biased value for some RBPs **(see Supplementary information)**.

In order classify genes as having or not having feedback (PABA) we need to decided on a threshold for the ability of introducing F as a free parameter to better explain the data (the increase in R^2^ upon varying F). To do so we fit the model to one biological replica of each gene, and calculated the R^2^ of that fit to data from an independent biological replica, so that the difference in R^2^ is an measure of biological and technical noise in the system. We found that the median of this differences was 0.0015, so we assumed that if the fit increases by more than this threshold value when allowing F_Tl_ to vary the feedback measurement is relevant (allowing three parameters to vary always increases the R^2^ of a fit compared to allowing two to vary). In addition, we find that 0.0015 is above the R^2^ increase in which the associated F diverges between positive and negative values **(Figure 3B)**, indicating that any gene that has a higher R^2^ increase has been measured to have a significant feedback strength. This reinforces this thresholding for assigning feedback. We note that these two methods are independent, and yet the agree on similar R^2^ increase threshold values.

### Mathematical modeling of the relationship between expression and fitness

Expression-vs-fitness data was modelled using a parametric impulse function (Chechik et al., 2008) based on two sigmoid functions fitted to the data. We fit the function as described by (Keren et al., 2016), in which seven free parameters define two sigmoid functions (to allow a proper modelling of both deleterious low and high expression) that predict the relationship between expression and fitness. We fit these to the measured expression-vs-fitness data (which constitutes the “presence of PABA” fit) and then inferred fitness from expression in the (predicted) absence of PABA (which corresponds to the “absence of PABA” fit). We next compared the GFP (promoter) - vs - fitness relationship in both the “presence” and “absence” models.

### Mathematical modeling of the limiting regulator model

The limiting regulator model was built considering that a fraction of steady state mRNA (calculated from equation (1), hereafter referred as initial [mRNA] (m_o_)) can be complexed with a Regulator (R), forming up an mRNA(m)-Regulator(R) complex (mR) as follows:

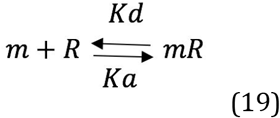

Where Ka and Kd are the reaction equilibrium constants. When any of the two factors can be limiting, the [mR] varies in time as:

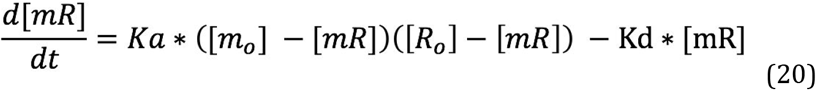

Where [m_o_] is the steady state [mRNA] described in eq (1) and R_o_ is the initial amount of regulator in the cell. Note that, at any given *dt, [m] = [m_o_] - [mR]* and *[R] = [R_o_] - [mR]*. At equilibrium, [mR] is:

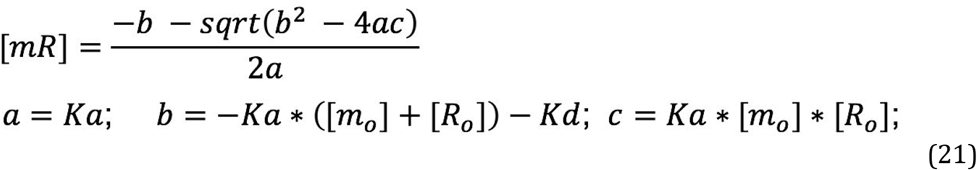

The uncomplexed remaining mRNA concentration ([m]) can be calculated as:

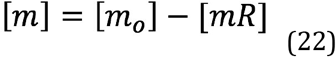

Then, protein concentrations ([RFP]) can be modelled as:

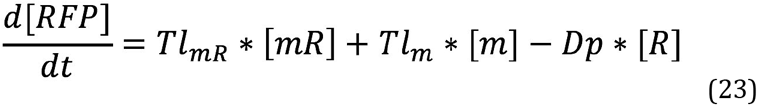

Where *Tl_mR_* is the translation rate of the mR complex, *Tl_m_* is the translation rate of [m], and *Dp* is the protein degradation rate. High *Tl_mR_* (compared to *Tl_m_*) corresponds to a positive regulator, and vice-versa for low *Tl_mR_*. At equilibrium, RFP concentrations correspond to:

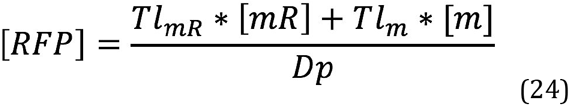

In order to model the *Z_3_EVpr-GFP vs Z_3_EVpr-mCherry-gene* profiles at different [R] we generated GFP and RFP data from equations (6) and (24), respectively. We changed the *Tl_m_* / *Tl_mR_* to generate positive and negative regulators, and varied *R_o_* to simulate several regulator concentrations in the cell (analogous to adding an extra chromosome set in a diploid).

## SUPPLEMENTARY INFORMATION

### SD (synthesis/degradation rates), BE_R_ (mCherry basal expression) and F (feedback rate) are the necessary free parameters for explaining the data

We previously described a similar system for measuring and modeling autoregulation(Schikora-Tamarit et al., 2016a). This initial framework assumed that any Z_3_EV_promoter_-mCherry-gene construct would alter mCherry intrinsic parameters such as mRNA or protein Synthesis or Degradation rates (hereafter referred as SD) and would not alter GFP expression or other mCherry features. Thus, the GFP induction curve should remain constant, and the mCherry induction curve should be offset by a constant amount (in log space). However, we found that both the GFP and the shape of the mCherry **(Figure 1)** induction curves vary across inducible the strains in the library, suggesting the need of a more complex mathematical framework for detecting PABA. One that uses GFP expression values as a control of the system’s induction.

This GFP and mCherry alterations lead to gene-specific mCherry vs GFP (RfG) profiles **(Figure S1)**. A visual examination of each of these profiles, compared to simulated curves in which each of the model parameters change **(Figure 2A)**, suggests a multiparametric alteration of BE_R_(as the fraction of maximum mCherry expression at zero induction levels), SD and/or F values in some of the constructs. Furthermore, a control mCherry construct (Y212) grown in sucrose and mannose (with a growth and metabolic difference to the dextrose-grown strain) showed a BE_R_-change-like behavior **(Figure S11)**. All these findings suggest that the model has to implement BE_R_ as a gene-specific parameter (which is not associated to PABA) for a proper quantification.

We investigated if BE_R_ can be calculated from experimental fluorescence data, in order to reduce the number of free parameters of the model. We found that our maximum induction level (100 nM) of the mCherry-gene doesn’t reach the maximum theoretical expression, as the predicted maximum mCherry from experimental data is always higher than the measured one **(Figure S12A)**, suggesting that BE_R_ can’t be calculated from fluorescence data. In addition, the predicted BE_R_ from a model fit is different from the measured BE_R_ for some RBPs **(Figure S12B)**. Therefore BE_R_has to be predicted as a free parameter. Surprisingly, the BE_R_ alterations are independent of the BE_G_ changes **(Figure S12C)**, suggesting that each fluorescent protein has it’s own basal induction rate, which can vary because of the specific construct.

Our model implements four SD parameters (transcription rate (Tc), mRNA degradation (Dm), translation rate (Tl) and protein degradation (Dp)) **(Figure S10A)**. Change in those four parameters results in identical changes to the RfG curve **(Figure S10B)**. Our experimental data cannot distinguish between each of these parameters. This suggests that any SD-like change of a mCherry-gene construct can be computed by any of these rates, so that we kept constant Tc_R_, Dm_R_ and Dp_R_ in our analysis to let Tl_R_ to be a free parameter proxy of SD rates.

Regarding PABA quantification, we considered four possible models, so that a gene can mediate a feedback loop through transcription (F_Tc_), translation (F_Tl_), mRNA degradation (F_Dm_) or protein degradation (F_Dp_). We found all of these four models behave the same across the range of simulated F values altering the RfG curve **(Figure S10C)**, so that we can’t distinguish at which step is each RBP modulating its own expression. We used the model in which the gene modulates translation efficiency (F_Tl_ as a feedback rate parameter) as a proxy for feedback (or PABA) strength. F ranges between −∞ and +∞ and is monotonic with positive or negative feedback strength (amplification or buffering, respectively).

All in all, Tl_R_, BE_R_ and F_Tl_ changes from a control (Y212) RfG curve give a set of parameters can explain all of RfG profiles measured in our library **(Figure S5)**.

### Fitness and metabolism changes don’t alter our ability to detect the PABA phenomena

One major assumption to build our system was that any change in metabolism, fitness and/or transcriptome (hereafter termed extrinsic factors) derived from an aberrant gene expression alters GFP and mCherry the same way, as they are driven by the same promoter. In order to test this we first measured GFP and mCherry control strain (Y212) grown in different carbon sources (SC + dextrose 2% (control), SC + dextrose 0.5%, SC + mannose 2%, and SC + sucrose 2%), in order to induce changes in those extrinsic factors. Growth rate was inferred from cell counts per second as measured by flow cytometry. We found that sucrose has a decreased fitness **(Figure S11A)**, and both GFP and mCherry induction curves are altered in mannose and sucrose **(Figures S11B,C)**, suggesting that metabolism and fitness alterations can change each of the fluorescent proteins’ expression.

Strikingly, this mCherry and GFP alteration is not proportional, as the GFP-vs-mCherry profile is not overlapping with the control strain for suc and man **(Figure S11D)**. However, the alteration is visually similar to a BE_R_ change **(Figure 2A)**, and a model fit to the suc and man data allowing BE_R_ to vary is enough to perfectly fit the data **(Figure S11E)**. As said above (see *SD (synthesis/degradation rates), BE_R_ (mCherry basal expression) and F (feedback rate) are the necessary free parameters for explaining the data*) this indicates that BE_R_-change-like effects can be due to alterations in extrinsic factors, so that we have to consider that any effect that can be explained by a BE_R_ is independent of feedback or PABA. Thus, implementing BE_R_ as a free parameter of our model allows us to overcome such artifacts.

On the other hand, we measured fitness of 12 library strains at different induction levels (see **Results**), which allowed us to assess if changes in growth rate secondary to gene aberrant expression are compromising any of our PABA measurements. We found that the control strain does not vary in fitness across *β*-estradiol induction levels. Therefore changes in expression of either GFP or mCherry do not impact fitness, and that the measured fitness effects are due to gene-specific, not mCherry, expression levels **(Figure S13A)**.

We also asked about the relationship between PABA and fitness vs expression profiles. A visual observation of the expression vs fitness results **(Figure S8)** indicates that each gene shows a characteristic fitness vs expression profile, allowing us its classification into fitness decrease at high expression (FDH), fitness decrease at low expression (FDL), non-monotonic fitness alteration (NMFA) and no fitness alteration (NFA). We can find genes with different PABA strength and sign across all of these categories **(Figures S8, Supplementary Table 1)**.

In summary, two lines of evidence suggest that variation in any extrinsic factor does not interfere with our feedback measurements: (1) genes exist with and without PABA and with and without fitness effects, and (2) if we experimentally alter fitness by changing the carbon source, the model does not require feedback to explain the experimental data.

### Supplementary Figures

**Figure S1.**
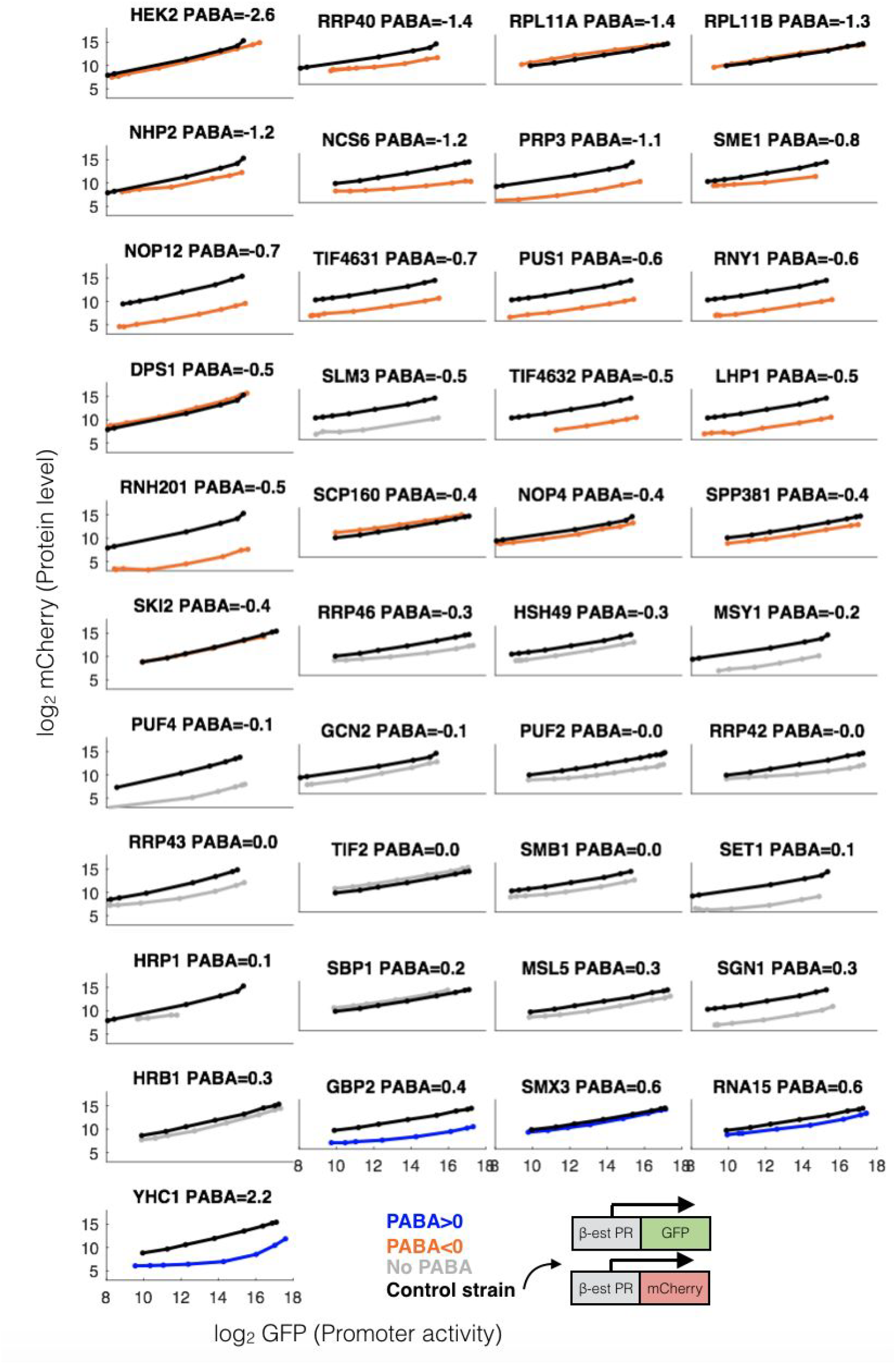
Each gene has a unique expression vs *β*-estradiol induction profile. Shown are log2 (mCherry) graphed against promoter activity (log2(GFP)).

**Figure S2.**
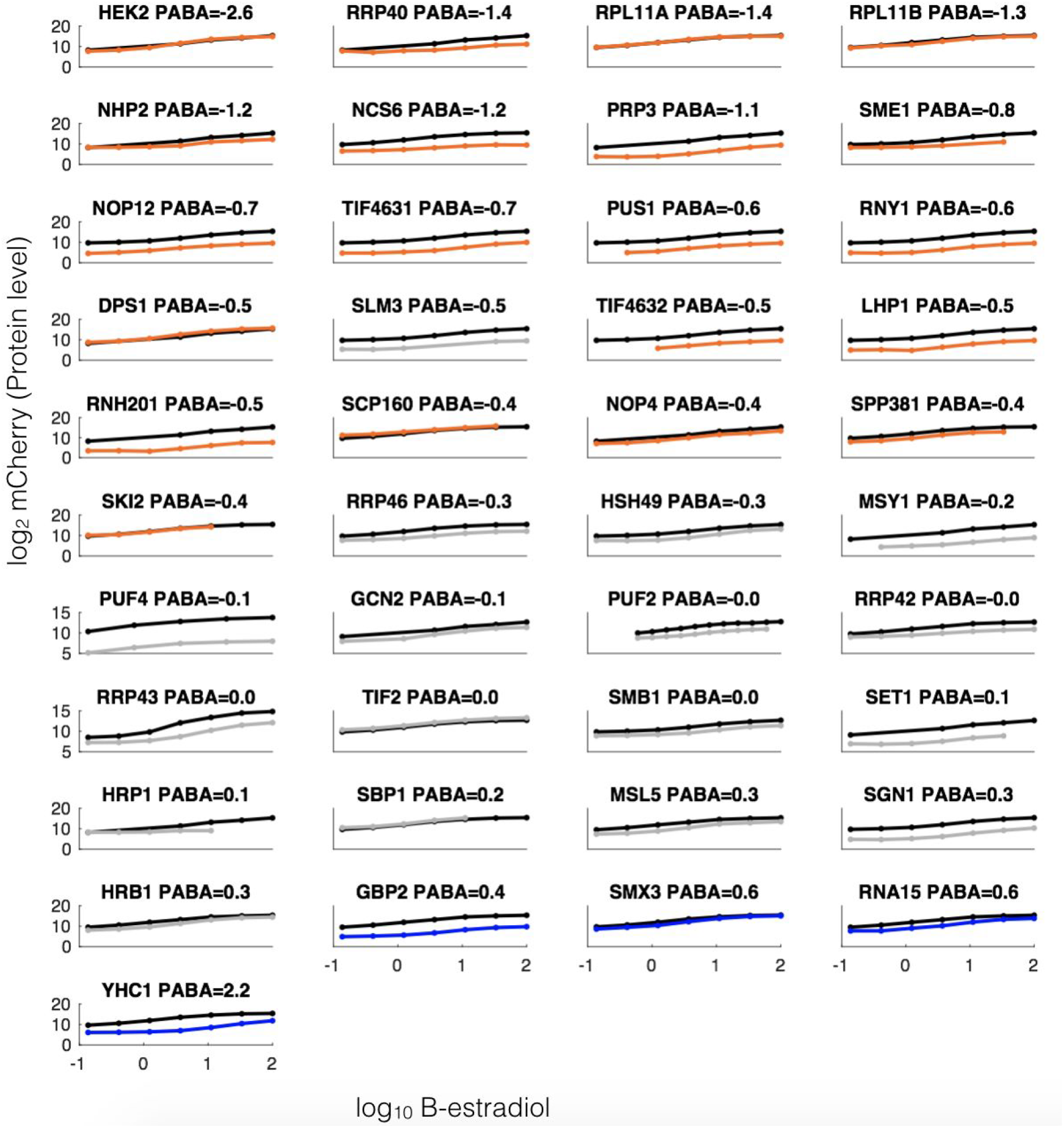
Each gene has a unique expression vs *β*-estradiol induction profile. Shown are log2 (mCherry) graphed against *β*-estradiol.

**Figure S3.**
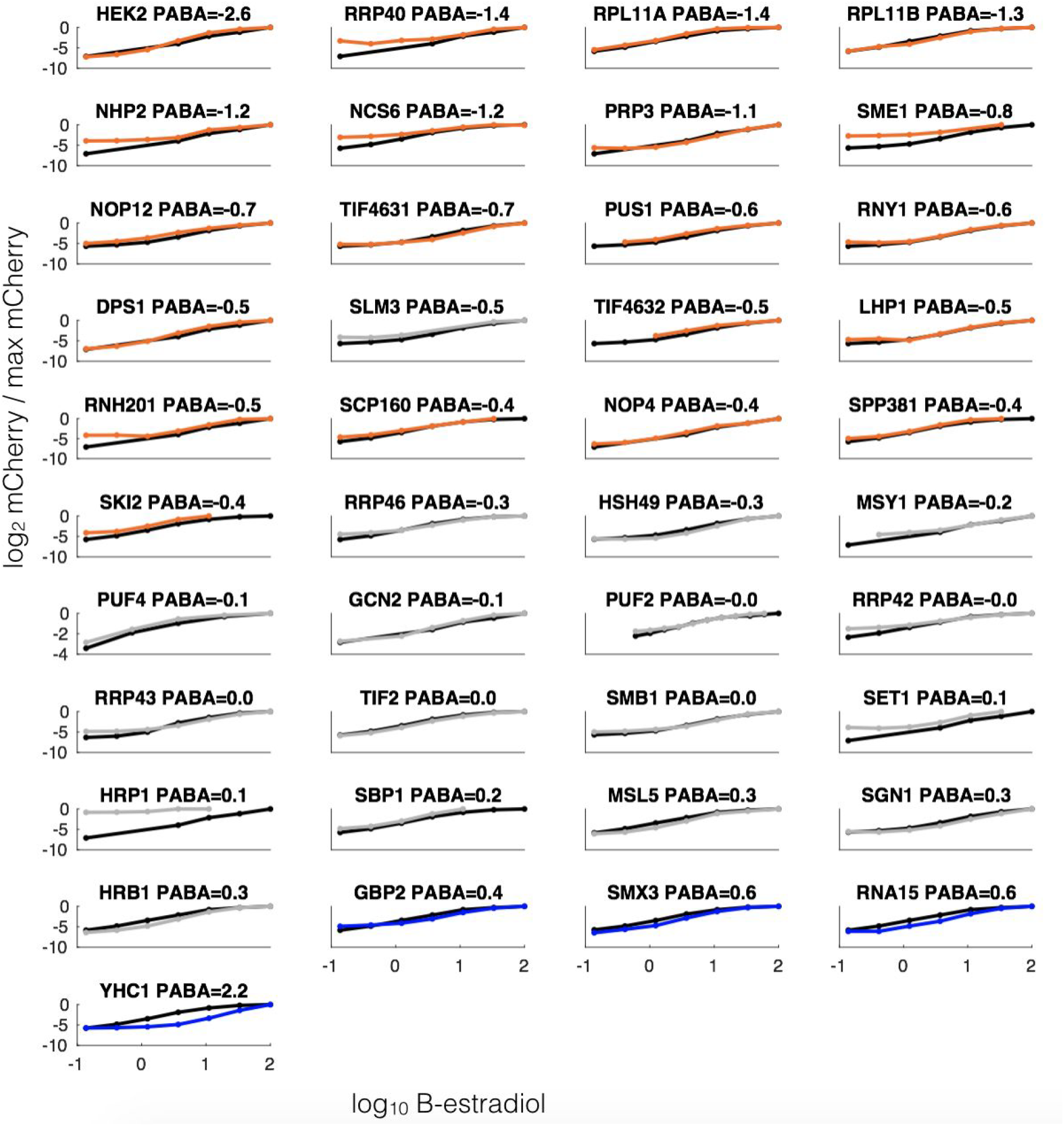
Each gene has a unique expression vs *β*-estradiol induction profile. All genes are shown with expression relative to that gene’s maximum mCherry expression.

**Figure S4.**
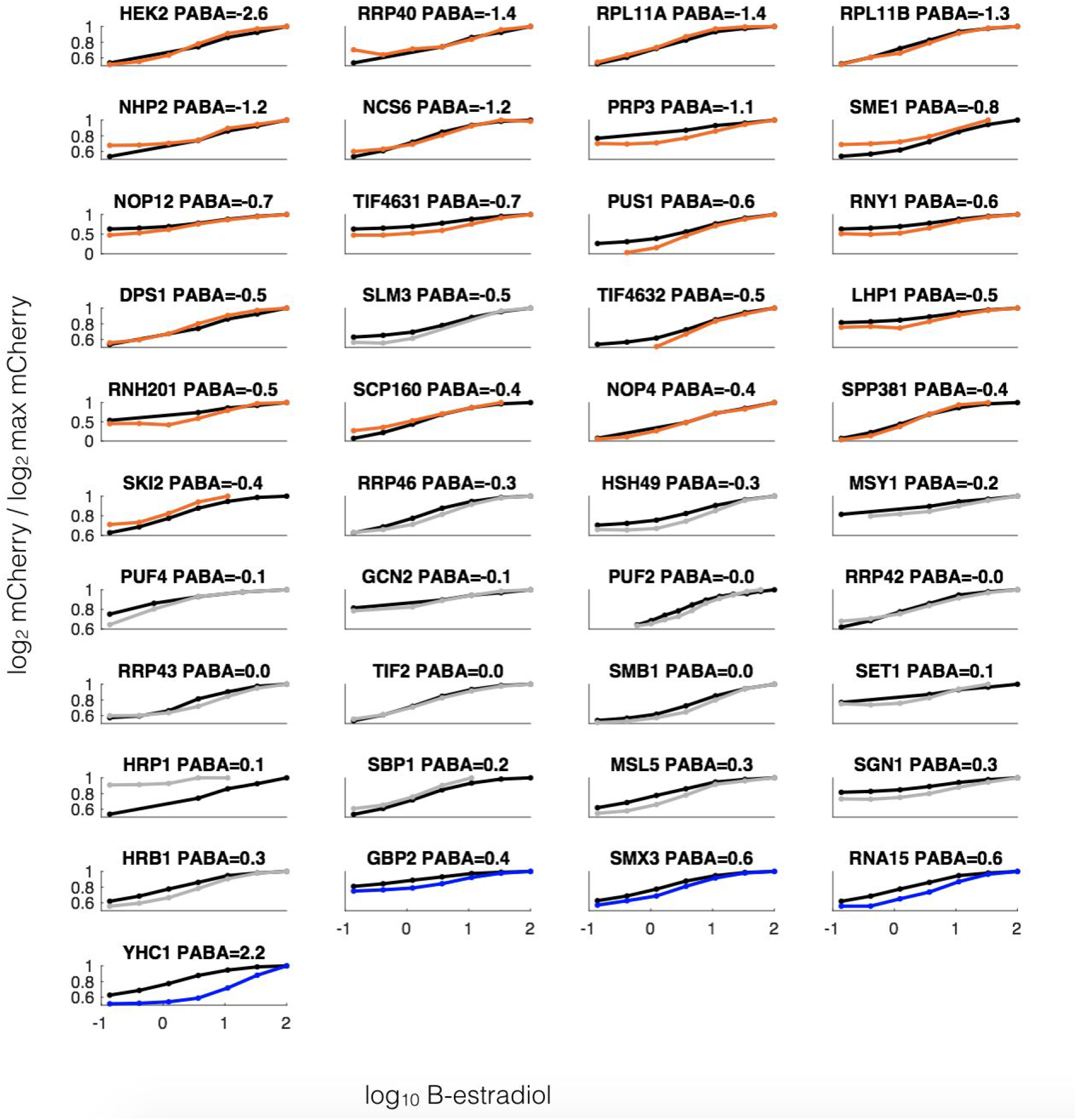
Each gene has a unique expression vs *β*-estradiol induction profile. All genes are shown with expression relative to that gene’s maximum mCherry expression.

**Figure S5.**
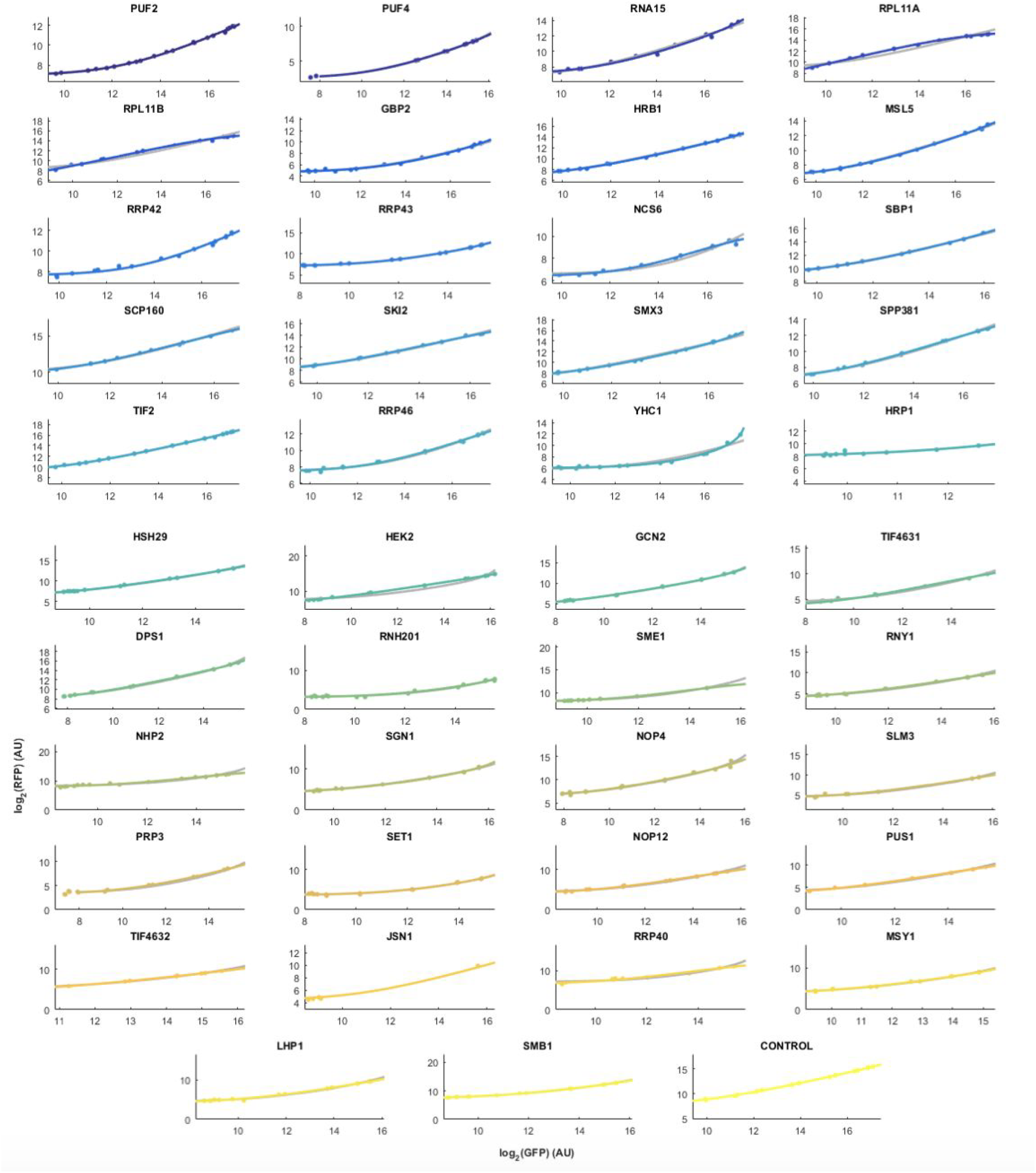
Model fits to the entire library. Shown is the log_2_ GFP vs log_2_ mCherry data (dots) fit by the model (lines), for each gene in the library.

**Figure S6.**
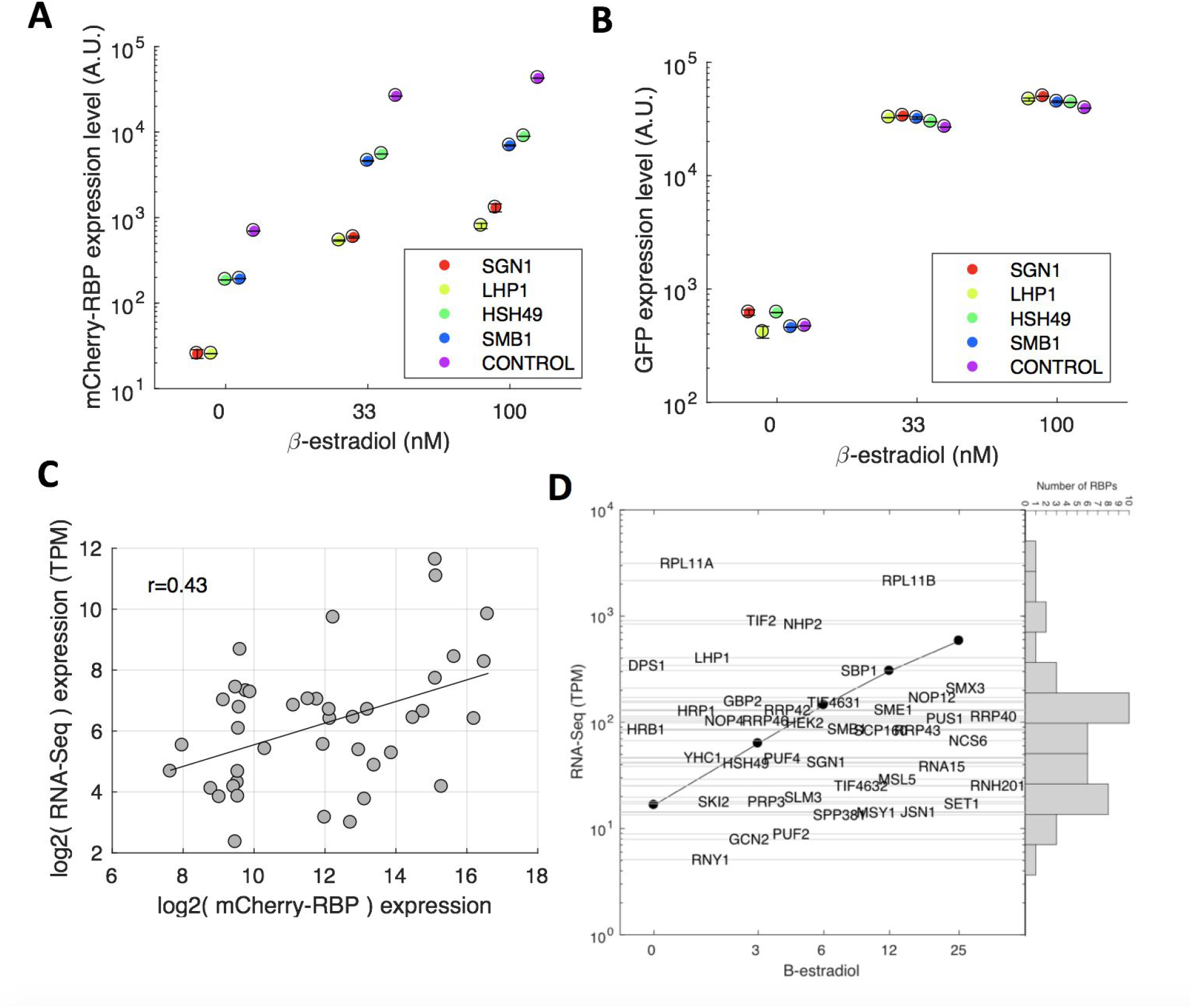
The dynamic range of induction for each gene includes native expression levels. **(A)** mCherry-gene expression levels vary over an order of magnitude in different tags across different induction levels. Shown is data from several example genes. **(B)** However, GFP expression is maintained in the same space. **(C)** mCherry at maximum induction is significantly correlated (p < 0.001) with measured native RNA-seq expression levels (taken from (Carey, 2015)). **(D)** RNA levels were calibrated to *β*-estradiol induction from a previous Z_3_EVpr-RPB9 and Z_3_EVpr-DST1 RNA-seq data (Carey, 2015), and are represented as black dots. Most of the gene’s native RNA expression overlap with this induction range, so that the measured feedback interactions are likely to be quantitatively determining native expression. The native TPM value for each gene is represented as the grey lines crossing each gene name, so that the X value for each gene name is random and does not correspond to *β*-estradiol. Shown is also the distribution of these RNA levels.

**Figure S7.**
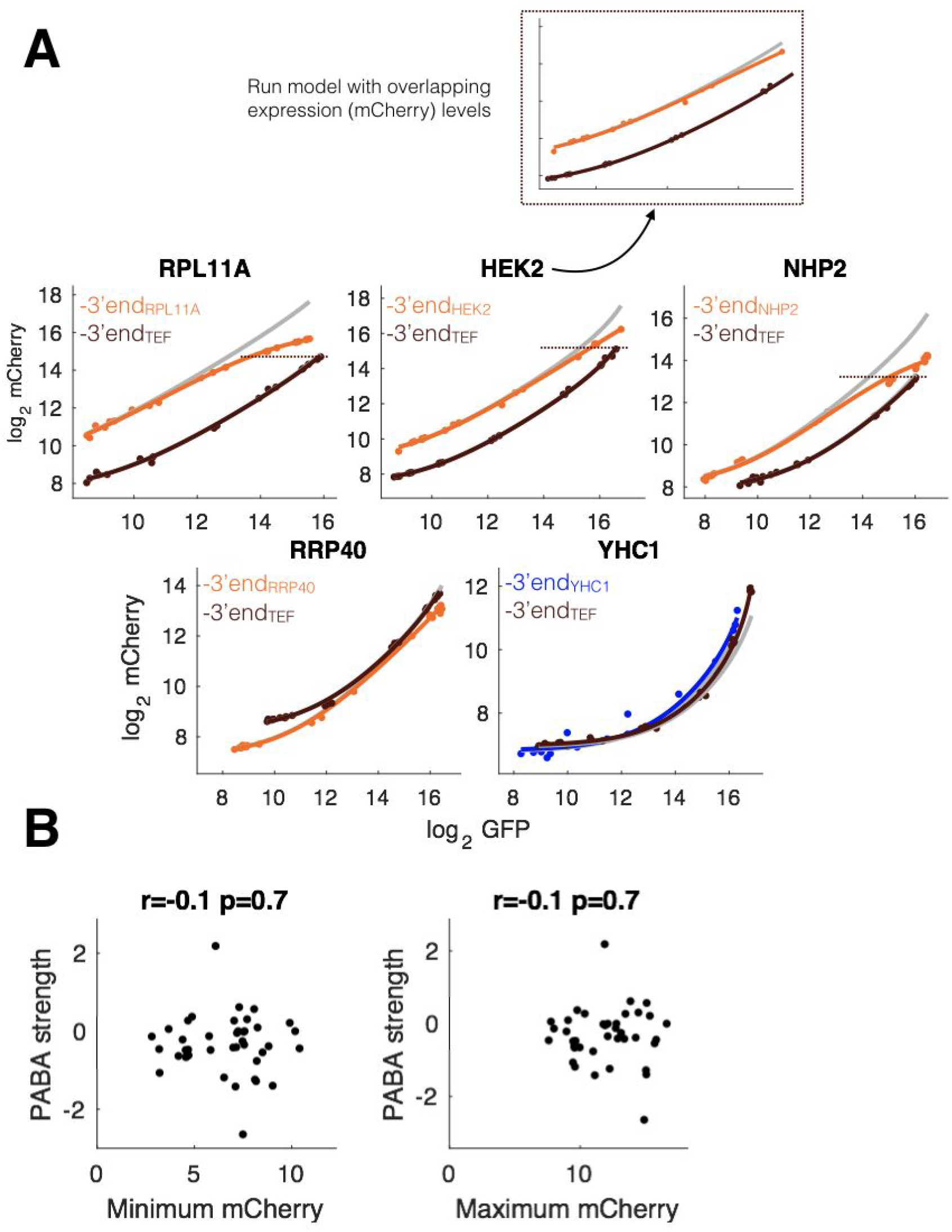
PABA can be encoded in 3’ end and ORF regions, and is not dependent on expression level. **(A)** Shown are the expression data for all genes for which we replaced the native 3’end with the TEF 3’end. The constructs and coloring correspond to those in **Figure 4C**. The orange and blue data and lines correspond to those constructs with the native 3’end. Brown lines and data refer to *mCherry-TEF_3’end_* constructs. The gray lines represent the absence of PABA. The upper panel represents the overlapping mCherry-protein range between native and TEF-3’end constructs. **(B)** PABA strength is not correlated with maximum (left) or minimum (right) mCherry-gene expression levels in our library.

**Figure S8.**
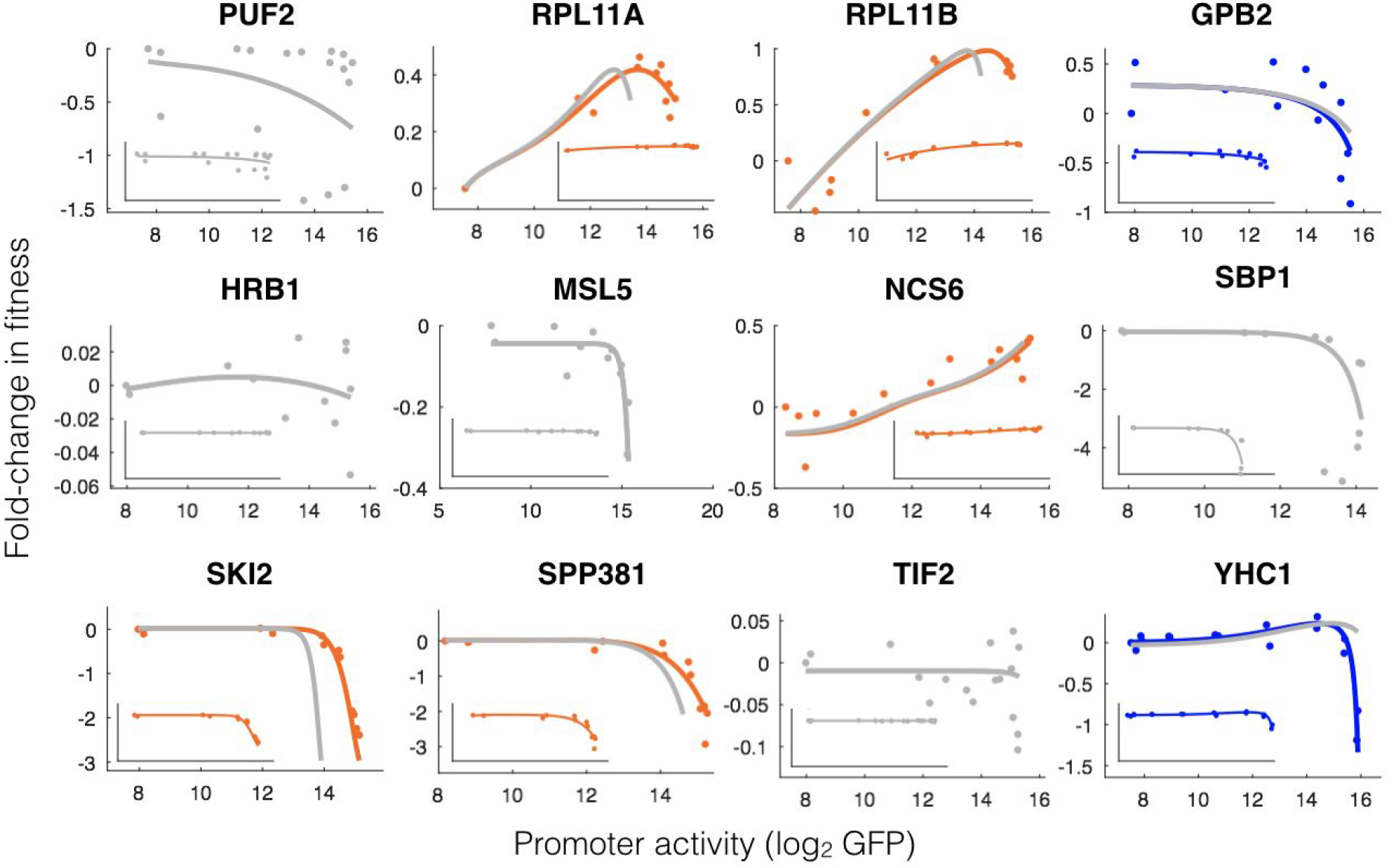
Each gene has a particular promoter-vs-fitness profile, affected by PABA. Fold-change in fitness (FCF) varies as a function of measured gene-mCherry expression for 9/12 genes for which fitness was measured. Two paralogs (RPL11A and RPL11B exhibit a non-monotonic relationship between expression and FCF, while 6/12 genes exhibit a decrease in FCF at high expression, and 1/12 (NCS6) exhibits an increase in FCF with increasing expression. The main panels have the y-axes adjusted to show all experimental variability within each gene. The insets all have the same y-axes so that variation in FCF can be compared across genes. The colored lines correspond to the model fit to the experimental data (dots) with PABA (orange for Buffering, blue for Amplification and gray for no PABA), while the gray lines correspond to the modelled absence of PABA.

**Figure S9:**
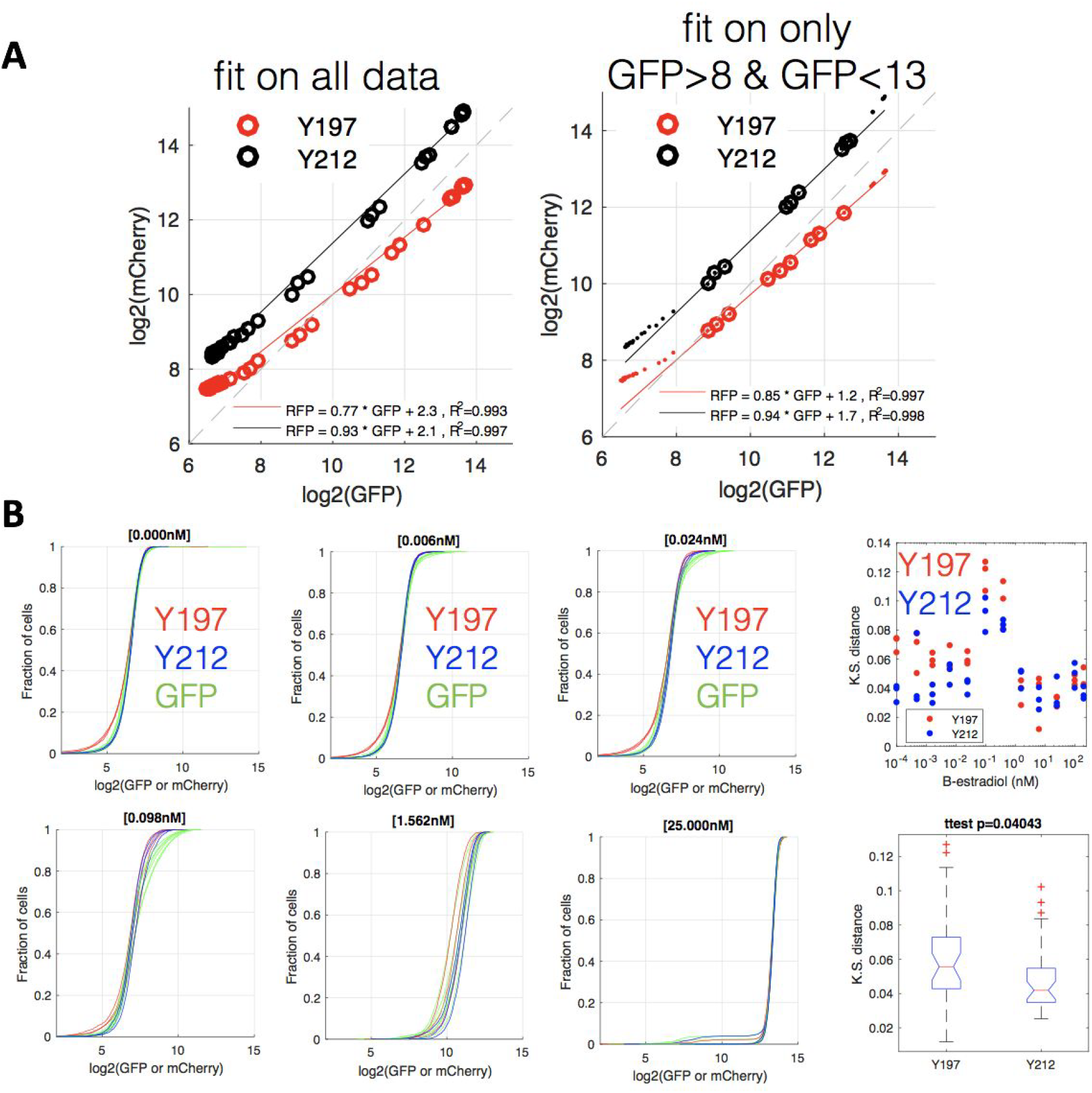
An AT-rich yeast codon optimized mCherry behaves more similarly to GFP at both the population level (across *β*-estradiol concentrations) and at the single cell level within a single *β*-estradiol concentration. **(A)** Shown are GFP vs mCherry for Y212 (his3::Z3EVpr-yco-mCherry) and Y197 (his3::Z3EVpr-mCherrτy). The Y212 data are more linear (higher R^2^ and slope closer to 1). Therefore the induction curves of the two fluors is more similar across *β*-estradiol concentrations. **(B)** Shown are the single-cell expression distributions for mCherry for Y197 and Y212, and GFP for Y212 (GFP is behaves identically between Y197 and Y212). Each line is one biological replicate. At low *β*-estradiol concentrations the yco-mCherry single-cell expression distribution is more similar to the GFP expression distribution (Kolmogorov–Smirnov test).

**Figure S10:**
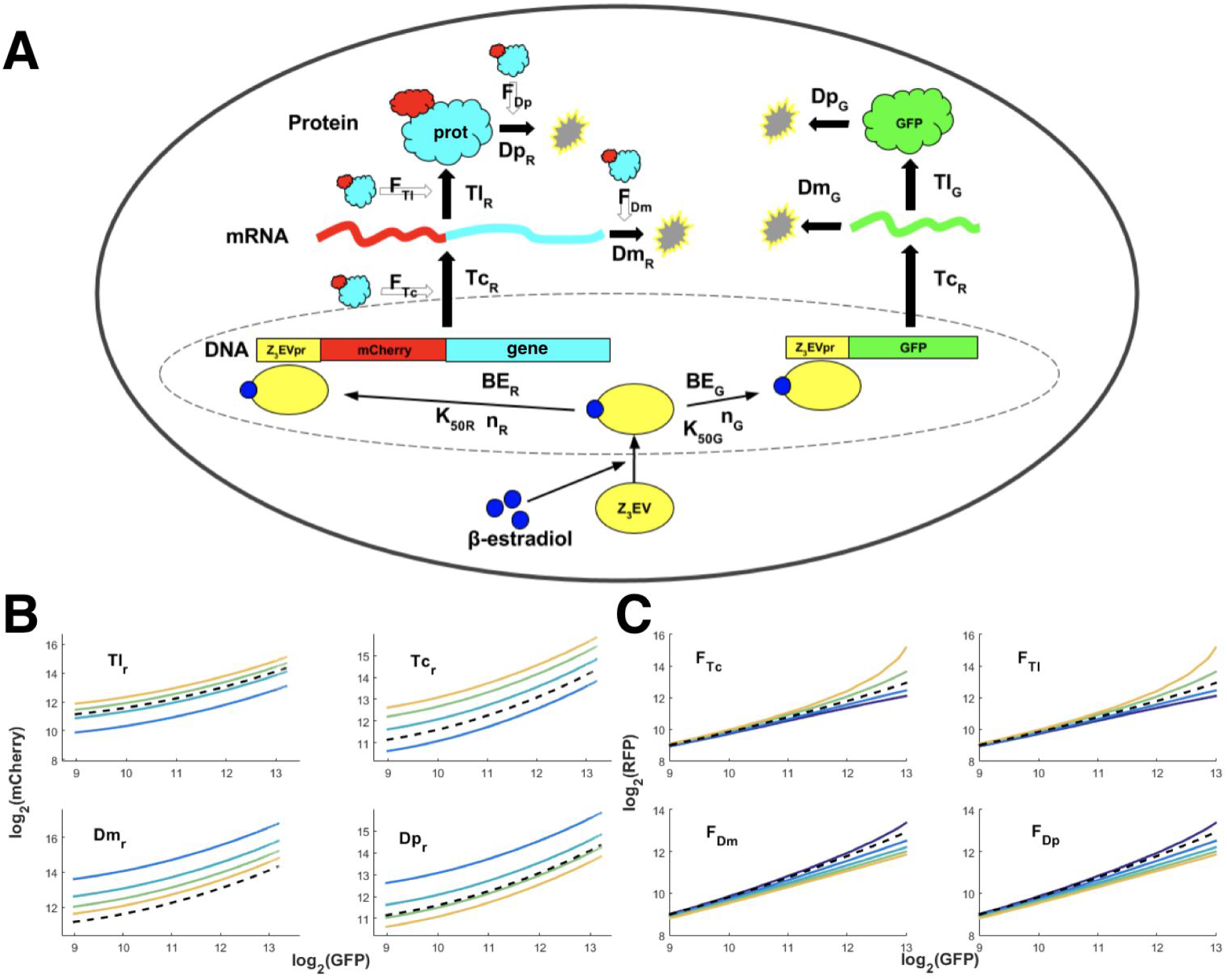
Variation in synthesis and degradation rates are experimentally indistinguishable in the model. **(A)** A simple model of gene expression, combined with our data analysis, predicts that GFP and mCherry-gene driven by the same Z3EVpr are going to be induced in a Hill-function manner with the same n (hill coefficient) and and constant K_50_ (*β*-estradiol concentration corresponding to half expression) ratio (see methods), whereas BE is going to be altered by each of the gene tags **(Figures S1, S2, S3, S4)**. Downstream of transcription start, each protein is expected to have it’s own transcription (Tc), translation (Tl), mRNA degradation (Dm) and protein degradation (Dp) rate. A feedback interaction, either direct or indirect, could be modulating Tc (F_Tc_), Tl(F_Tl_), Dm (F_Dm_) or Dp (FD_Dp_) so that each of the F would correspond to a value between −∞ and +∞ monotonic with positive or negative feedback strength, respectively. **(B)** Our model can’t distinguish between each of the **4** parameters that conform the synthesis/degradation rates (SD), as changes in any of them (colored lines) depict the same offset-like behavior in terms of a RfG simulation, relative to a reference value (black dashed). **(C)** Any of the 4 models mentioned in (A) depict a similar GFP-vs-mCherry profile for different values of F (colored lines) relative to a reference (black dashed), with matching behavior among synthesis-mediated feedback (upper) and degradation-mediated feedback (lower).

**Figure S11:**
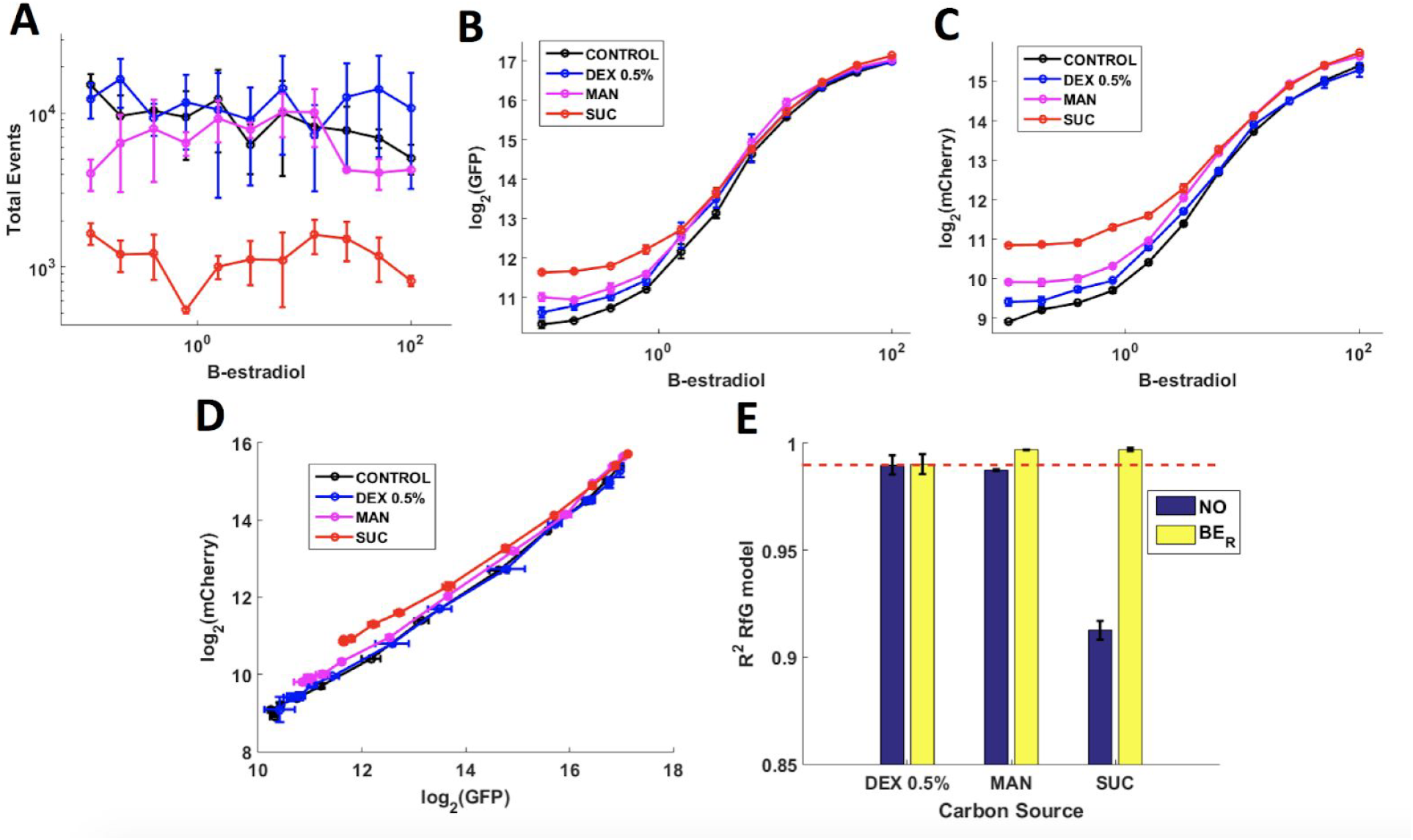
Metabolism and fitness alter BE_R_. **(A)** A control strain grown in SC with different carbon sources (Dextrose (Dex) 2% as control, Dex 0.5 %, Mannose (Man) 2 % and Sucrose (Suc) 2 %) and *β*-estradiol was assayed. Growth rate (inferred from total FACS event count) is impaired in sucrose-based media, but not altered in mannose and dextrose 0.5%. **(B,C)** Each growth condition alters both the GFP (B) and mCherry (C) induction curves, and sucrose data shows an mCherry BE-increase-like behavior not coupled to a GFP BE proportional increase.**(D)** The GFP-vs-mCherry profile visually shows an mCherry BE change-like behavior (see **Figure 2A**) in Man and Suc, whereas Dex 0.5 % depicts the same profile as control. **(E)** The Man and Suc data (specially Suc) can be explained by a change in mCherry BE, as the R^2^ increases giving a perfect fit when allowing it to vary as a free parameter (yellow) relative to the case in which the Dex 2% set of parameters are fit (blue), something that doesn’t happen with the Dex 0.5% data. The red dashed line refers to the R^2^ got from the Dex 0.5% data which has a good fit to the control parameters, so that’s useful as a threshold for good fit. Error bars refer to the SEM of the different replicates’ fit.

**Figure S12:**
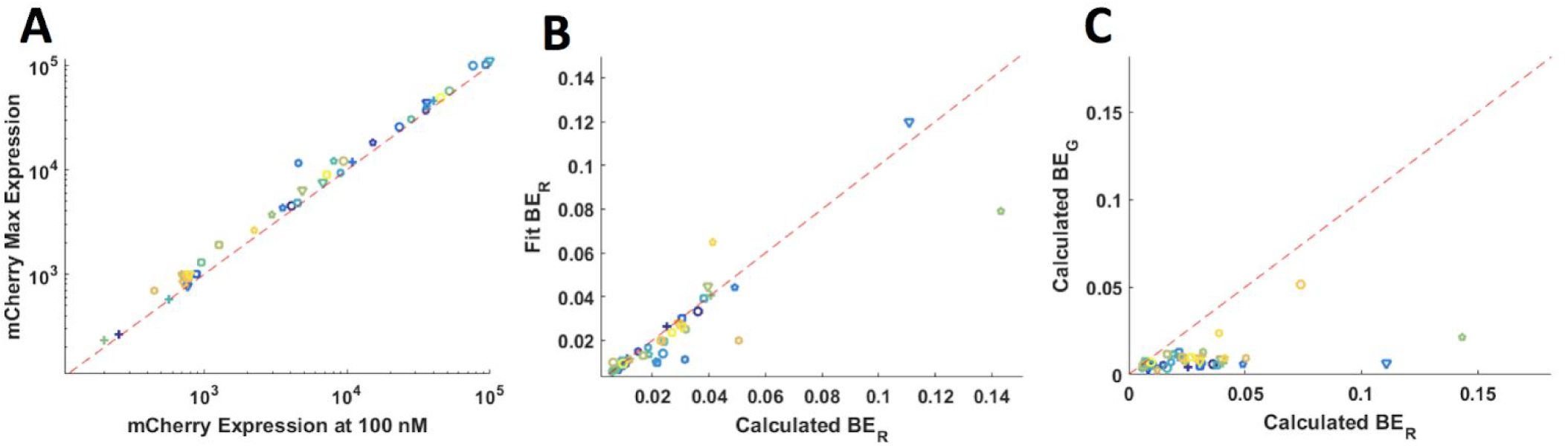
Each gene alters the mCherry basal expression (BE_R_) in a non-measurable way. **(A)** A model fit mCherry data predicts that the system doesn’t reach maximum expression at 100 nM *β*-estradiol, so that we can’t measure max expression for some of the constructs. **(B)** Calculating BE from measured fluorescence data gives a slightly different value from the one got of an mCherry data fit for some of the constructs. **(C)** Each of the genes alters the mCherry BE (BE_R_) in an uncoupled GFP BE (BE_G_) way.

**Figure S13:**
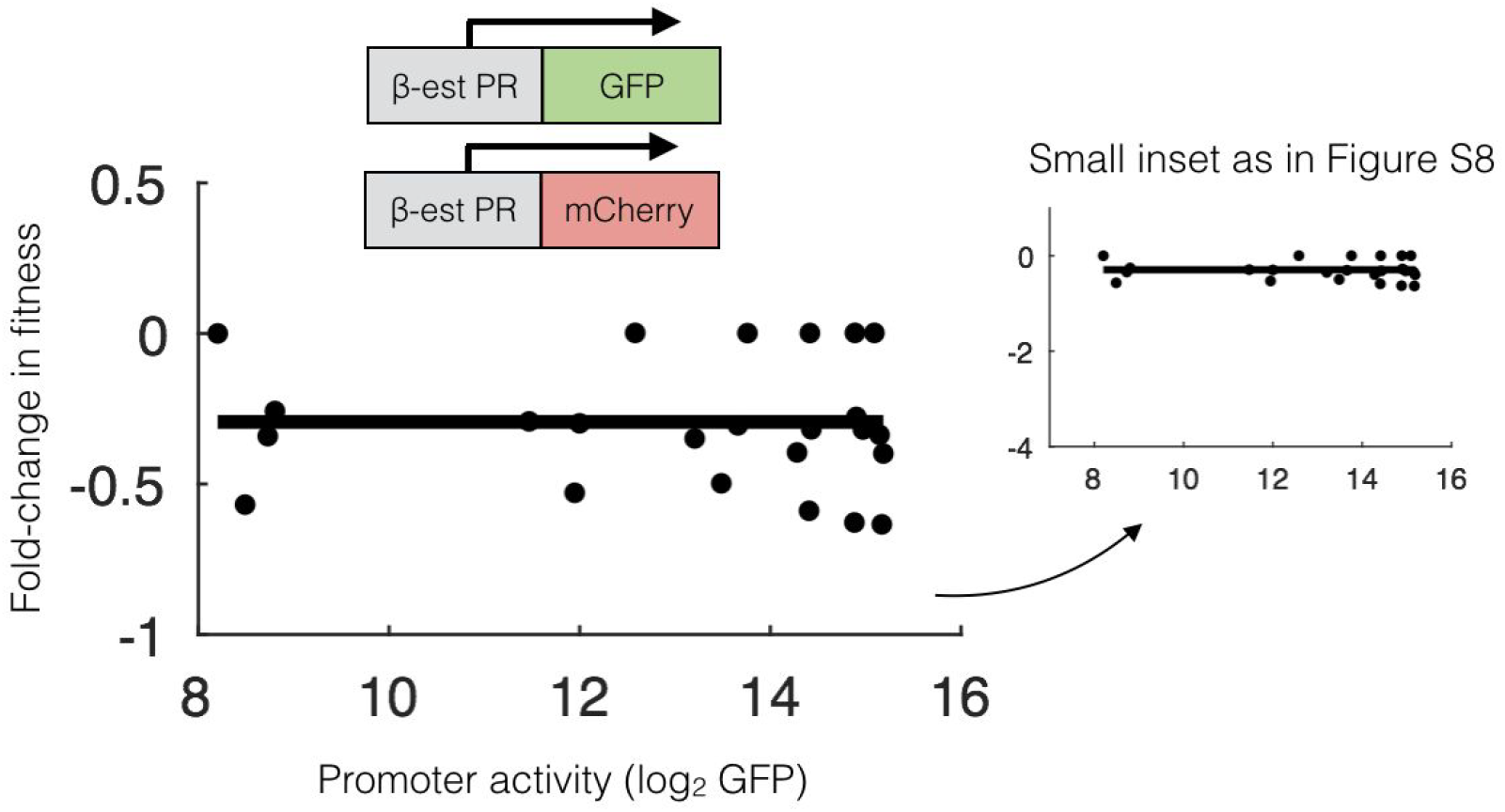
Fitness does not vary as a function of mCherry in the control strain. **(A)** Shown is the data similar to **Figure 8D** and **Figure S8** for the fluorescence control strain.

**Figure S14:**
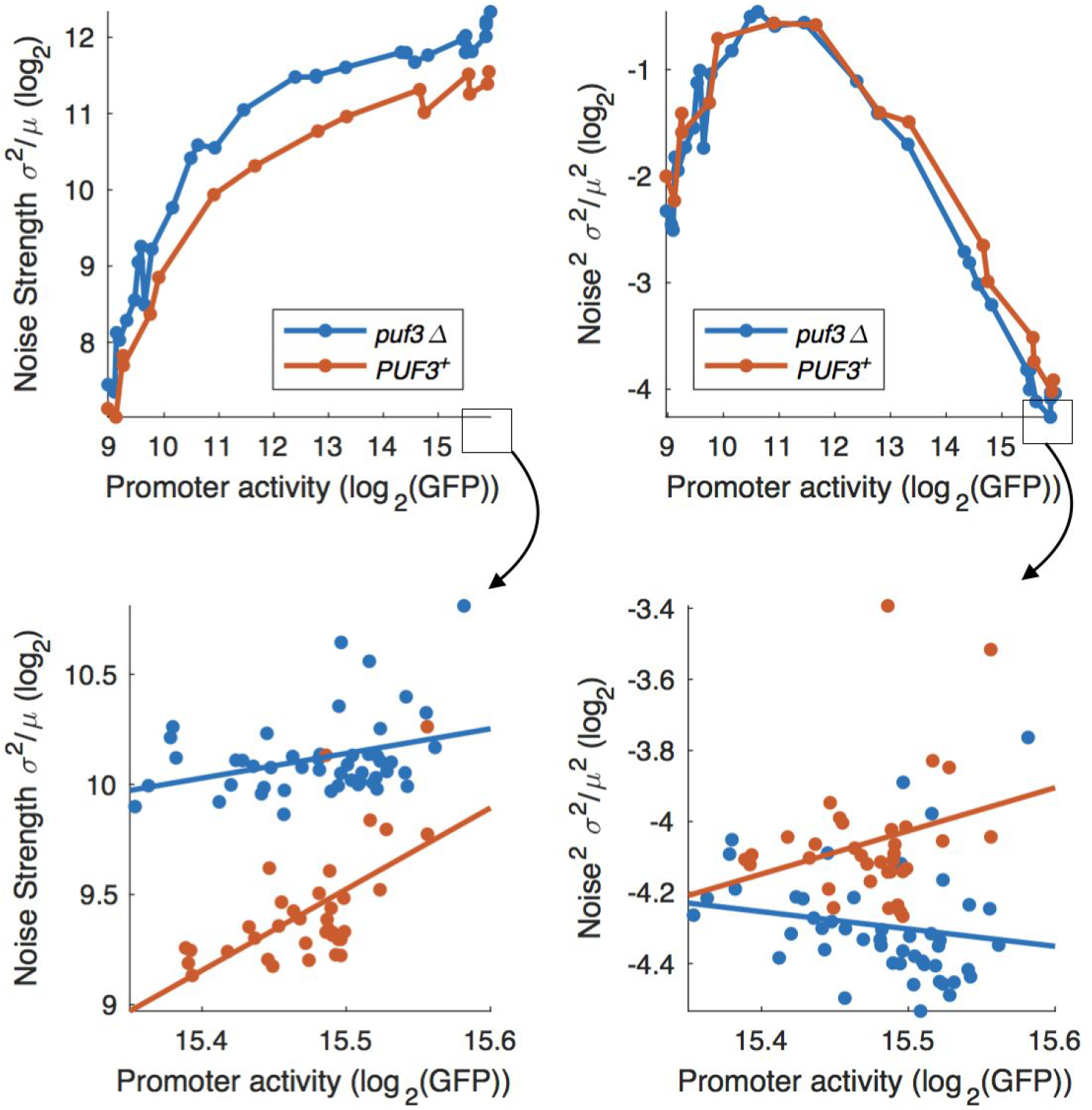
Changes in noise strength for the mCherry-COX17_3’end_ strain are consistent with a change in mRNA stability and suggest titration of limiting Puf3p at high *β*-estradiol. Shown are total Noise^2^ and Noise Strength for the mCherry-COX17_3’end_ construct grown at low (0-200nM, top) and high (62 - 2000nM, bottom). Noise strength is expected to increase in a model in which Puf3 decreases the mCherry-COX17 mRNA stability(Paulsson, 2005). Higher noise strength (Fano factor) in the *puf3* strain is consistent with this model. Interestingly, noise strength increases at very high [*β*-estradiol] in the PUF3^+^ strain, but not the *puf3* strain. This suggests that high expression of *mCherry-COX17_3’end_* may titrate limiting Puf3 protein.

**Figure S15.**
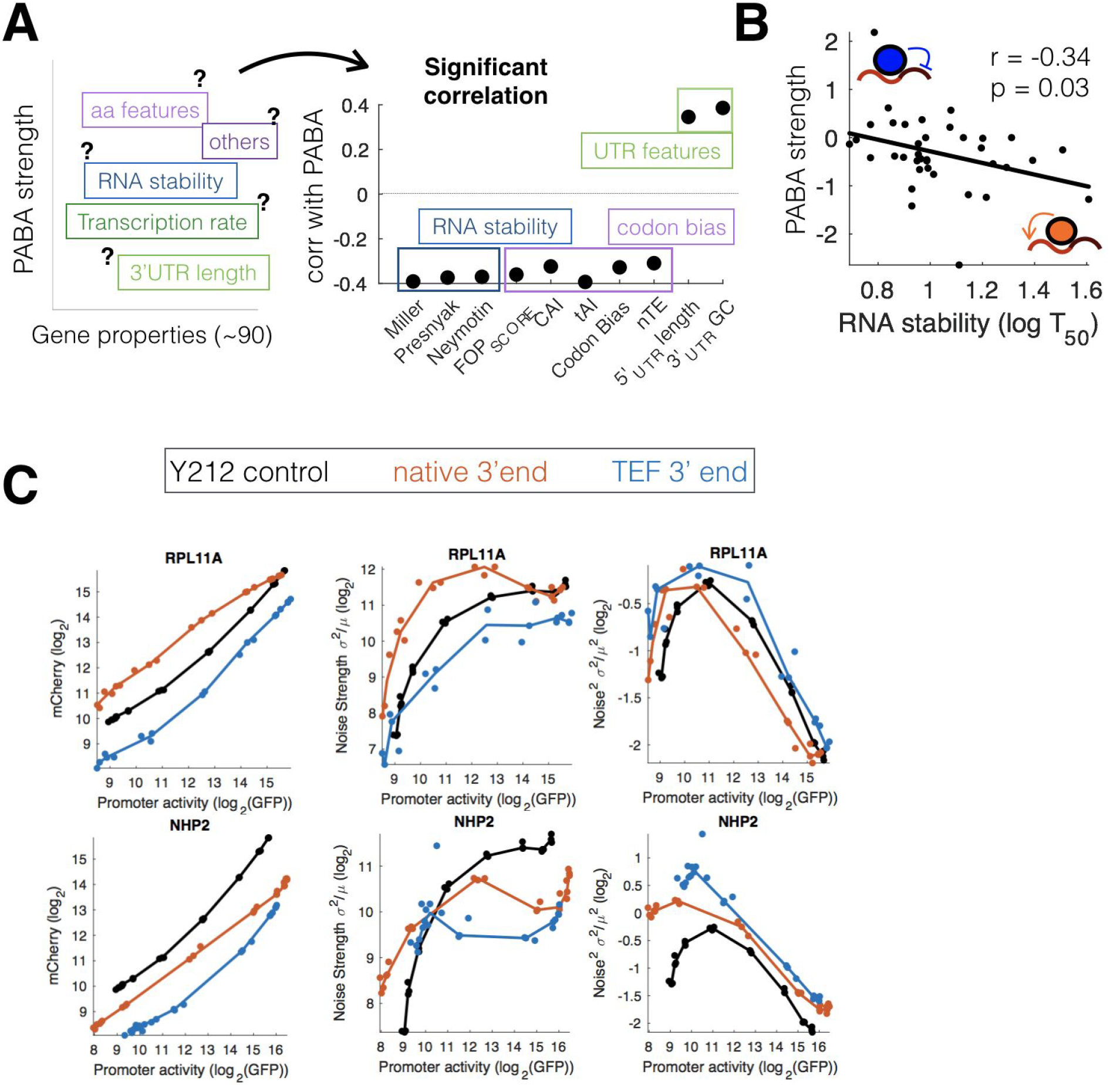
PABA is partially associated with mRNA stability. **(A)** We searched a set of 89 different gene properties (such as transcription rate, mRNA stability) for features that correlate with PABA. **(B)** Shown are the only features that significantly correlate with PABA strength, grouped by biological meaning. PABA strength is negatively correlated with mRNA stability and sequences features related to mRNA stability. **(C)** Shown are noise and noise strength graphed against GFP for mCherry-gene, for the RPL11A and NHP2 constructs. RPL11A with the native 3’end exhibits a decrease in noise strength at high [TF], suggesting a decrease in translation efficiency or mRNA or protein stability. RPL11A-TEF_3’end_ exhibits no PABA, and no such decrease. Such conclusions cannot be drawn for the NHP2 constructs.

**Figure S16.**
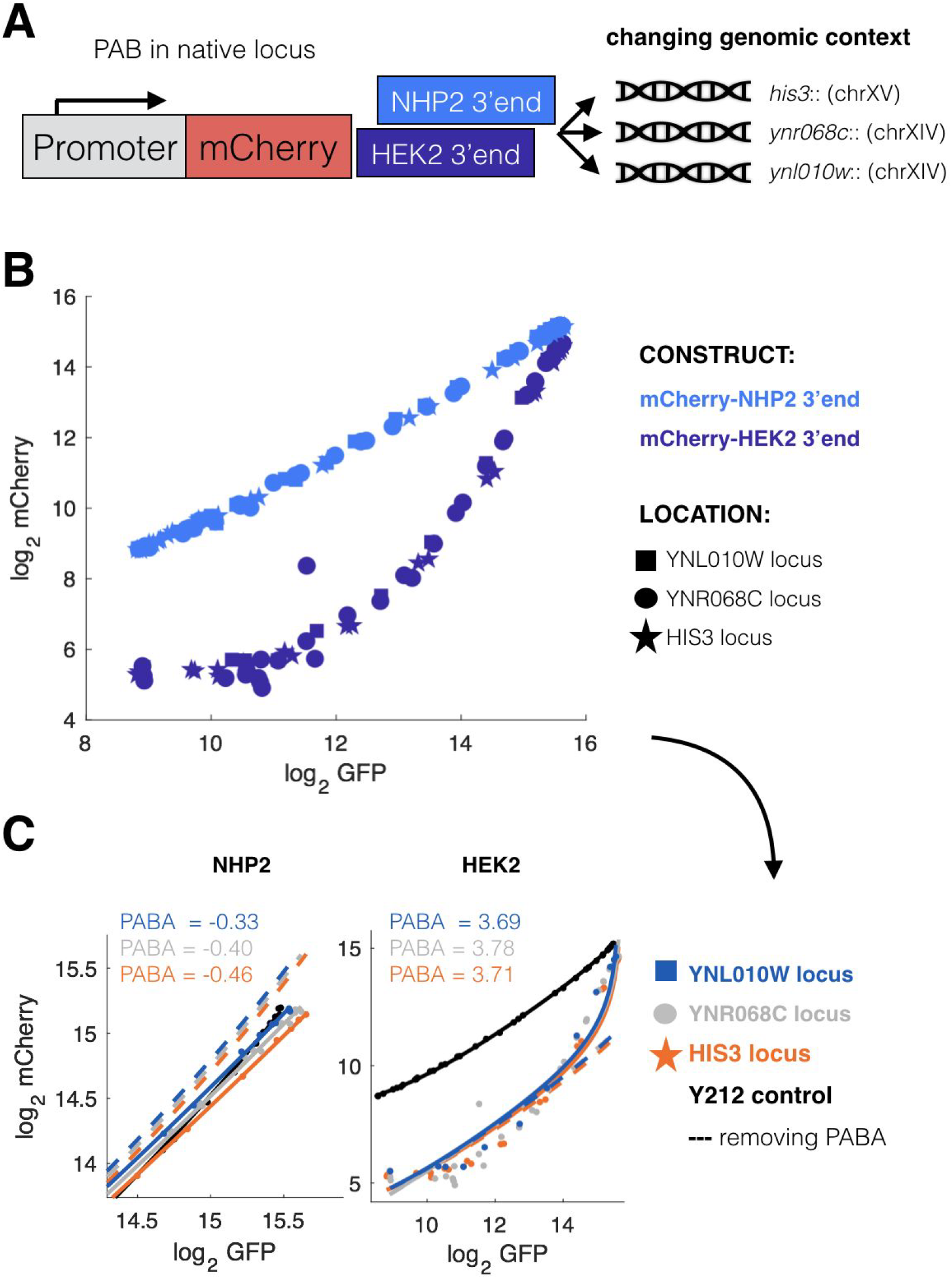
PABA is independent of genomic context. **(A)** We integrated *Z_3_EVpr-mCherry-3’end* constructs for NHP2 and HEK2 (with strong native possibly 3’end-mediated PAB) at three different loci: HIS3 and two positions in chromosome XIV. **(B)** The promoter-vs-expression profile clusters by construct (colors) rather than genomic position (markers). **(C)** All mCherry-3’end_NHP2_ constructs have PAB, while the mCherry-3’end_HEK2_ have PAA. The full lines represent a model fit with PABA, and the dashed ones without PABA, as in **Figure 2F**.

## REFERENCES

Albert, F.W., Kruglyak, L., 2015. The role of regulatory variation in complex traits and disease. Nat. Rev. Genet. 16, 197–212.

Ang, J., Harris, E., Hussey, B.J., Kil, R., McMillen, D.R., 2013. Tuning response curves for synthetic biology. ACS Synth. Biol. 2, 547–567.

Baudrimont, A., Voegeli, S., Viloria, E.C., Stritt, F., Lenon, M., Wada, T., Jaquet, V., Becskei, A., 2017. Multiplexed gene control reveals rapid mRNA turnover. Sci Adv 3, e1700006.

Becskei, A., Serrano, L., 2000. Engineering stability in gene networks by autoregulation. Nature 405, 590–593.

Brachmann, C.B., Davies, A., Cost, G.J., Caputo, E., Li, J., Hieter, P., Boeke, J.D., 1998. Designer deletion strains derived from Saccharomyces cerevisiae S288C: a useful set of strains and plasmids for PCR-mediated gene disruption and other applications. Yeast 14, 115–132.

Brewster, R.C., Weinert, F.M., Garcia, H.G., Song, D., Rydenfelt, M., Phillips, R., 2014. The transcription factor titration effect dictates level of gene expression. Cell 156, 1312–1323.

Brophy, J.A.N., Voigt, C.A., 2014. Principles of genetic circuit design. Nat. Methods 11, 508–520.

Buchler, N.E., Louis, M., 2008. Molecular titration and ultrasensitivity in regulatory networks. J. Mol. Biol. 384, 1106–1119.

Carey, L.B., 2015. RNA polymerase errors cause splicing defects and can be regulated by differential expression of RNA polymerase subunits. Elife 4. https://doi.org/10.7554/eLife.09945

Carey, L.B., van Dijk, D., Sloot, P.M.A., Kaandorp, J.A., Segal, E., 2013. Promoter sequence determines the relationship between expression level and noise. PLoS Biol. 11, e1001528.

Chechik, G., Oh, E., Rando, O., Weissman, J., Regev, A., Koller, D., 2008. Activity motifs reveal principles of timing in transcriptional control of the yeast metabolic network. Nat. Biotechnol. 26, 1251–1259.

Chen, S., Li, K., Cao, W., Wang, J., Zhao, T., Huan, Q., Yang, Y.-F., Wu, S., Qian, W., 2017. Codon-Resolution Analysis Reveals a Direct and Context-Dependent Impact of Individual Synonymous Mutations on mRNA Level. Mol. Biol. Evol. 34, 2944–2958.

Dvir, S., Velten, L., Sharon, E., Zeevi, D., Carey, L.B., Weinberger, A., Segal, E., 2013. Deciphering the rules by which 5’-UTR sequences affect protein expression in yeast. Proc. Natl. Acad. Sci. U. S. A. 110, E2792–801.

Eriksson, P.R., Ganguli, D., Nagarajavel, V., Clark, D.J., 2012. Regulation of histone gene expression in budding yeast. Genetics 191, 7–20.

Espinar, L., Schikora Tamarit, M.À., Domingo, J., Carey, L.B., 2018. Promoter architecture determines cotranslational regulation of mRNA. Genome Res. https://doi.org/10.1101/gr.230458.117

Gietz, R.D., Woods, R.A., 2006. Yeast transformation by the LiAc/SS Carrier DNA/PEG method. Methods Mol. Biol. 313, 107–120.

Gonçalves, E., Fragoulis, A., Garcia-Alonso, L., Cramer, T., Saez-Rodriguez, J., Beltrao, P., 2017. Widespread Post-transcriptional Attenuation of Genomic Copy-Number Variation in Cancer. Cell Syst 5, 386–398.e4.

Hansen, A.S., O’Shea, E.K., 2015. cis Determinants of Promoter Threshold and Activation Timescale. Cell Rep. 12, 1226–1233.

Harigaya, Y., Parker, R., 2016. Codon optimality and mRNA decay. Cell Res. 26, 1269–1270.

Ishikawa, K., Makanae, K., Iwasaki, S., Ingolia, N.T., Moriya, H., 2017. Post-Translational Dosage Compensation Buffers Genetic Perturbations to Stoichiometry of Protein Complexes. PLoS Genet. 13, e1006554.

Kaplan, N., Moore, I.K., Fondufe-Mittendorf, Y., Gossett, A.J., Tillo, D., Field, Y., LeProust, E.M., Hughes, T.R., Lieb, J.D., Widom, J., Segal, E., 2009. The DNA-encoded nucleosome organization of a eukaryotic genome. Nature 458, 362–366.

Karr, J.R., Sanghvi, J.C., Macklin, D.N., Gutschow, M.V., Jacobs, J.M., Bolival, B., Jr, Assad-Garcia, N., Glass, J.I., Covert, M.W., 2012. A whole-cell computational model predicts phenotype from genotype. Cell 150, 389–401.

Keren, L., Hausser, J., Lotan-Pompan, M., Vainberg Slutskin, I., Alisar, H., Kaminski, S., Weinberger, A., Alon, U., Milo, R., Segal, E., 2016. Massively Parallel Interrogation of the Effects of Gene Expression Levels on Fitness. Cell. https://doi.org/10.1016/j.cell.2016.07.024

Keren, L., Zackay, O., Lotan-Pompan, M., Barenholz, U., Dekel, E., Sasson, V., Aidelberg, G., Bren, A., Zeevi, D., Weinberger, A., Alon, U., Milo, R., Segal, E., 2013. Promoters maintain their relative activity levels under different growth conditions. Mol. Syst. Biol. 9, 701.

Kim, E., Hart, T., 2017. All for One, and One for All. Cell Syst 5, 314–316.

Kim, H.D., O’Shea, E.K., 2008. A quantitative model of transcription factor-activated gene expression. Nat. Struct. Mol. Biol. 15, 1192–1198.

Kimura, M., 1983. The Neutral Theory of Molecular Evolution. Cambridge University Press.

Kintaka, R., Makanae, K., Moriya, H., 2016. Cellular growth defects triggered by an overload of protein localization processes. Sci. Rep. 6, 31774.

Levo, M., Segal, E., 2014. In pursuit of design principles of regulatory sequences. Nat. Rev. Genet. 15, 453–468.

López-Maury, L., Marguerat, S., Bähler, J., 2008. Tuning gene expression to changing environments: from rapid responses to evolutionary adaptation. Nat. Rev. Genet. 9, 583–593.

McIsaac, R.S., Oakes, B.L., Wang, X., Dummit, K.A., Botstein, D., Noyes, M.B., 2013. Synthetic gene expression perturbation systems with rapid, tunable, single-gene specificity in yeast. Nucleic Acids Res. 41, e57–e57.

McShane, E., Sin, C., Zauber, H., Wells, J.N., Donnelly, N., Wang, X., Hou, J., Chen, W., Storchova, Z., Marsh, J.A., Valleriani, A., Selbach, M., 2016. Kinetic Analysis of Protein Stability Reveals Age-Dependent Degradation. Cell 167, 803–815.e21.

Mnaimneh, S., Davierwala, A.P., Haynes, J., Moffat, J., Peng, W.-T., Zhang, W., Yang, X., Pootoolal, J., Chua, G., Lopez, A., Trochesset, M., Morse, D., Krogan, N.J., Hiley, S.L., Li, Z., Morris, Q., Grigull, J., Mitsakakis, N., Roberts, C.J., Greenblatt, J.F., Boone, C., Kaiser, C.A., Andrews, B.J., Hughes, T.R., 2004. Exploration of essential gene functions via titratable promoter alleles. Cell 118, 31–44.

Nevozhay, D., Adams, R.M., Murphy, K.F., Josić, K., Balázsi, G., 2009. Negative autoregulation linearizes the dose–response and suppresses the heterogeneity of gene expression. Proceedings of the National Academy of Sciences. https://doi.org/10.1073/pnas.0809901106

Olivas, W., Parker, R., 2000. The Puf3 protein is a transcript-specific regulator of mRNA degradation in yeast. EMBO J. 19, 6602–6611.

Paulsson, J., 2005. Models of stochastic gene expression. Phys. Life Rev. 2, 157–175.

Phillips, R., 2015. Theory in Biology: Figure 1 or Figure 7? Trends Cell Biol. https://doi.org/10.1016/j.tcb.2015.10.007

Plotkin, J.B., Kudla, G., 2011. Synonymous but not the same: the causes and consequences of codon bias. Nat. Rev. Genet. 12, 32–42.

Radhakrishnan, A., Chen, Y.-H., Martin, S., Alhusaini, N., Green, R., Coller, J., 2016. The DEAD-Box Protein Dhh1p Couples mRNA Decay and Translation by Monitoring Codon Optimality. Cell 167, 122–132.e9.

Rest, J.S., Morales, C.M., Waldron, J.B., Opulente, D.A., Fisher, J., Moon, S., Bullaughey, K., Carey, L.B., Dedousis, D., 2013. Nonlinear fitness consequences of variation in expression level of a eukaryotic gene. Mol. Biol. Evol. 30, 448–456.

Ryan, C.J., Kennedy, S., Bajrami, I., Matallanas, D., Lord, C.J., 2017. A Compendium of Co-regulated Protein Complexes in Breast Cancer Reveals Collateral Loss Events. Cell Syst 5, 399–409.e5.

Schikora-Tamarit, M.À., Toscano-Ochoa, C., Domingo Espinós, J., Espinar, L., Carey, L.B., 2016a. A synthetic gene circuit for measuring autoregulatory feedback control. Integr. Biol. 8, 546–555.

Schikora-Tamarit, M.À., Toscano-Ochoa, C., Espinós, J.D., Espinar, L., Carey, L.B., 2016b. A synthetic gene circuit for measuring autoregulatory feedback control. Integr. Biol. https://doi.org/10.1039/C5IB00230C

Segal, E., Widom, J., 2009. From DNA sequence to transcriptional behaviour: a quantitative approach. Nat. Rev. Genet. 10, 443–456.

Shalem, O., Carey, L., Zeevi, D., Sharon, E., Keren, L., Weinberger, A., Dahan, O., Pilpel, Y., Segal, E., 2013. Measurements of the Impact of 3′ End Sequences on Gene Expression Reveal Wide Range and Sequence Dependent Effects. PLoS Comput. Biol. 9, e1002934.

Shalem, O., Sharon, E., Lubliner, S., Regev, I., Lotan-Pompan, M., Yakhini, Z., Segal, E., 2015. Systematic dissection of the sequence determinants of gene 3′ end mediated expression control. PLoS Genet. 11, e1005147.

Shoval, O., Alon, U., 2010. SnapShot: network motifs. Cell 143, 326–e1.

Sopko, R., Huang, D., Preston, N., Chua, G., Papp, B., Kafadar, K., Snyder, M., Oliver, S.G., Cyert, M., Hughes, T.R., Boone, C., Andrews, B., 2006. Mapping pathways and phenotypes by systematic gene overexpression. Mol. Cell 21, 319–330.

Springer, M., Weissman, J.S., Kirschner, M.W., 2010. A general lack of compensation for gene dosage in yeast. Mol. Syst. Biol. 6, 368.

Sung, M.-K., Porras-Yakushi, T.R., Reitsma, J.M., Huber, F.M., Sweredoski, M.J., Hoelz, A., Hess, S., Deshaies, R.J., 2016. A conserved quality-control pathway that mediates degradation of unassembled ribosomal proteins. Elife 5. https://doi.org/10.7554/eLife.19105

Thattai, M., 2016. Universal Poisson Statistics of mRNAs with Complex Decay Pathways. Biophys. J. 110, 301–305.

Tsai, I.J., Bensasson, D., Burt, A., Koufopanou, V., 2008. Population genomics of the wild yeast Saccharomyces paradoxus: Quantifying the life cycle. Proc. Natl. Acad. Sci. U. S. A. 105, 4957–4962.

Uwimana, N., Collin, P., Jeronimo, C., Haibe-Kains, B., Robert, F., 2017. Bidirectional terminators in Saccharomyces cerevisiae prevent cryptic transcription from invading neighboring genes. Nucleic Acids Res. 45, 6417–6426.

van Dijk, D., Sharon, E., Lotan-Pompan, M., Weinberger, A., Carey, L., Segal, E., 2015. Competition between binding sites determines gene expression at low transcription factor concentrations. bioRxiv. https://doi.org/10.1101/033753

van Dijk, D., Sharon, E., Lotan-Pompan, M., Weinberger, A., Segal, E., Carey, L.B., 2017. Large-scale mapping of gene regulatory logic reveals context-dependent repression by transcriptional activators. Genome Res. 27, 87–94.

Wagner, A., 2005. Energy constraints on the evolution of gene expression. Mol. Biol. Evol. 22, 1365–1374.

Yamanishi, M., Ito, Y., Kintaka, R., Imamura, C., Katahira, S., Ikeuchi, A., Moriya, H., Matsuyama, T., 2013. A genome-wide activity assessment of terminator regions in Saccharomyces cerevisiae provides a “terminatome” toolbox. ACS Synth. Biol. 2, 337–347.

Yuan, Y., Guo, L., Shen, L., Liu, J.S., 2007. Predicting gene expression from sequence: a reexamination. PLoS Comput. Biol. 3, e243.

Zeevi, D., Lubliner, S., Lotan-Pompan, M., Hodis, E., Vesterman, R., Weinberger, A., Segal, E., 2014. Molecular dissection of the genetic mechanisms that underlie expression conservation in orthologous yeast ribosomal promoters. Genome Res. 24, 1991–1999.

Zenklusen, D., Larson, D.R., Singer, R.H., 2008. Single-RNA counting reveals alternative modes of gene expression in yeast. Nat. Struct. Mol. Biol. 15, 1263–1271.

